# Uncovering key genes related to potato tuber starch concentration from the tuber bulking, maturation stage to storage

**DOI:** 10.64898/2026.07.28.741397

**Authors:** Yu Chenhui, Yu Xiaogang, Jiang Bo, Liu Zhiru, Li Hui, Ao Xiang, Wang Jingshun, Qiu Ping, Zhang Lingkui, Bai Jianming, Shi Ying, Li Jieping

**Affiliations:** College of Agriculture, Northeast Agricultural University, Harbin 150030, China); Asia-Pacific Center, International Potato Center (China), Beijing 100200, China); Hulunbuir Academy of Agricultural and Animal Husbandry Sciences, Hailar District, Hulunbuir City, Inner Mongolia Autonomous Region 021000, China); Chinese Academy of Agricultural Sciences, Beijing, 100081, China); Yunnan Academy of Agricultural Sciences, Yunnan, 650051, China)

**Keywords:** Potato, High-low starch potato population, Starch and sucrose metabolic pathways, Transcriptome analysis, Three stages, β-glucosidase

## Abstract

Potato is the world’s fourth major staple crop, and tuber starch is a critical dietary energy source and a food processing raw material. In this study, hybrid progenies of the high-starch cultivar Huasheng No.7 and China main commercial varieties were used as materials, high-throughput sequencing was performed on tuber samples at tuber bulking, maturation and storage stages. Combined with bioinformatics analysis, parent-progeny resequencing, SNP screening, the regulatory mechanism of tuber starch metabolism was explored. The results showed significant stage-specific transcriptional reprogramming in potato tubers: few differentially expressed genes (DEGs) were identified at tuber bulking and maturation stages, but massive transcriptional changes occurred during storage. XET family genes regulated cell wall remodeling and carbon translocation across all stages, and WRKY transcription factors specifically controlled starch homeostasis in stored tubers. The DEGs at different stage were induced by the SNPs from their parent, and the allele from the high-starch parent, Huasheng 7 may contribute the high-starch allele to the offsprings. One key gene, *Soltu.DM.02G019910,* which encoded β-glucosidase, have a negatively relationship with tuber starch concentration. The low-starch potato breeding tubers shown higher significantly β-glucosidase activity than high-starch breeding population at maturation stage. This study clarifies the molecular regulatory network of potato tuber starch metabolism and screens core regulatory genes from the tuber bulking stage to tuber storage, thereby providing theoretical support and genetic resources for molecular breeding of high-starch and storage-resistant potato varieties.

## 1 Introduction

Potato starch, a natural polymer compound with predominantly round or oval-shaped granules, exhibits superior processing characteristics compared to starches from other crops. Its gel properties, transparency, water retention capacity, and oil retention ability surpass those of corn and wheat starches (Waterschoot *et al*., 2015). Breeders and researchers had developed high starch potato varieities and identified high starch potato. Zhao *et al*. analyzed 14 potato cultivars, revealing a starch content range of 13.07%-19.13% with a coefficient of variation of 12.2%. Varieties such as Dutch 2-12, Mengshu 17, and Jizhangshu 18 demonstrated notably high starch content exceeding 17% (Zhao *et al*., 2011). Bai *et al*. found cultivar had the largest influence the starch yield, and Yunshu 201 reached the highest starch yield of 8.16 t·hm□² (Bai *et al*., 2006). Yunshu 202 (late-maturing variety) with an average tuber starch content of 18.8% based on multi-year measurements (Zhang *et al*., 2019; Li *et al*., 2020; Li *et al*., 2018); Chuanyu 16 (early-maturing high-starch variety) with 23.9% starch content (Li *et al*., 2018), and Dongnong 308 with over 18% starch content (Li *et al*., 2017). After decades of long-term breeding efforts, the Potato Research Institute of Hulunbuir Academy of Agricultural and Animal Sciences has developed a series of high-starch potato varieties, including Velas, Huasheng No.7, and Neishu No.7. These varieties consistently exhibit starch content exceeding 18% over multiple years (Liu, 1999; Jiang *et al*., 2020; Jiang *et al*., 2024).

Some key starch regulation relative genes were uncovered and already be utilized in improving potato starch traits. Adegbaju *et al*. reduced expression of potato SBE1 and GWD1 genes via RNA interference (RNAi), revealing that these two genes collectively govern starch chain length distribution, starch particle morphology, and gelatinization viscosity characteristics (Adegbaju *et al*., 2025). Using potato Clearwater Russet as experimental material, researchers found that *PTST2b* maintains normal heteromorphic growth of starch particles; silencing *PTST2b* alters particle morphology and increases amylose content, whereas overexpression of the *PTST2*-interacting MRC gene enhances starch particle initiation sites, induces particle deformities, and reduces tuber yield and starch accumulation (Seung *et al*., 2015;Seung *et al*., 2018; Hochmuth *et al*., 2025). Hu *et al*. employing the diploid potato AC142, edited the *SS5* gene via CRISPR/Cas9 and demonstrated that *SS5* mutations reduce chloroplast starch accumulation, alter tuber starch particle size and number, decrease tuber fresh weight and starch content, increase amylose-to-globular fraction ratio, and lower gelatinization temperature, confirming *SS5* as a pivotal gene regulating potato starch particle formation (Hu *et al*., 2025). Singh *et al*. edited the *ESV1* and *LESV* genes in potatoes, demonstrating that *ESV1* and *LESV* participate in starch degradation by regulating the activity of GWD and PWD kinases, thereby jointly governing the glucan structure on starch granule surfaces and enzymatic hydrolysis efficiency (Singh *et al*., 2022). Locquet *et al*. further found that LESV gene mutations significantly reduce starch granule size, enhance starch phosphorylation levels and hydrolysis efficiency, and alter starch microstructure, whereas *ESV1* mutations have minimal effects on starch degradation-related phenotypes (Locquet *et al*., 2026). Sharma *et al*. utilized CRISPR/Cas9 technology to edit the *Pho1a* gene in the Desiree potato strain, obtaining multiple mutant lines that showed no impact on plant growth but increased tuber number, reduced single-tuber weight and amylose content, and altered starch granule size distribution (Sharma *et al*., 2023). Using the low-temperature saccharification-sensitive potato variety E3, Liu *et al*. silenced the *StTST1* gene via RNA interference, revealing that this silencing decreased reducing sugar and acrylamide levels post-low temperature storage while improving processing quality without affecting normal plant growth (Liu *et al*., 2023). Silencing the *StTST3.1* gene was shown to regulate transient starch turnover and chlorophyll synthesis in potato leaves; mutations in this gene induced plant dwarfism and yellowing while also modulating expression of key enzymes involved in starch synthesis and degradation (Liu *et al*., 2023). Ai *et al*. conducted experiments using the potato Desiree cultivar and found that the AP2/ERF transcription factor *StERF75* negatively regulates potato starch synthesis by inhibiting the expression of the *ISA1.1* gene, thereby reducing amylopectin content and altering starch chain length distribution and gelatinization properties (Ai *et al*., 2025). The research team led by Li demonstrated that the potato NAC-f transcription factor *StNAC033* suppresses the expression of genes involved in starch synthesis and degradation, thereby inhibiting the degradation of transient nighttime starch in potatoes (Li, 2024). Relevant studies have confirmed that the WRKY family IIc transcription factor *StWRKY23* directly targets the StSBE2.1 gene, regulating starch synthesis through epigenetic modifications; silencing this gene significantly increases potato tuber starch content and inhibits tuber bulking (<u>Sun, 2023</u>). Cui research revealed that the heat shock transcription factor *StHSFA2* activates key starch-degrading genes such as StBAM1 and *StUGPase2*, promoting the conversion of starch to reducing sugars and influencing the quality of potato storage and processing under low temperatures (Cui, 2025). Liu research showed that the ABA signaling pathway transcription factor *ScAREB2* binds to and upregulates the expression of the *ScBAM3.1* gene, accelerating starch granule degradation and markedly increasing soluble sugar content in potato tubers (Liu, 2023). The Larichev research team utilized CRISPR/Cas9 technology to edit the GBSS gene in potato varieties Nevsky and Udacha, generating genetically modified mutant materials that demonstrated the gene’s involvement in regulating starch synthesis in potatoes (Larichev *et al*., 2022). The Manabayeva team performed CRISPR/Cas9 editing on the GBSS gene in three potato varieties—Astanalyk, Tokhtar, and Aksor—and found that mutants with varying genetic backgrounds all exhibited a significant reduction in starch content (Abeuova *et al*., 2023). Ali *et al*. edited the GBSS gene in the potato strain Desiree to obtain various allele-edited lines, and confirmed through iodine staining that loss of GBSS function significantly reduces starch content in potato tubers (Ali *et al*., 2023). Jayarathna *et al*. knocked out all four GBSS alleles in Desiree material, revealing a substantial decrease in starch content and confirming GBSS as the core gene for starch synthesis in potatoes (Jayarathna *et al*., 2024). Zhao *et al*. edited the SBEI and SBEII genes in Desiree, demonstrating that multi-site editing of both genes significantly increased starch content, eliminated amylopectin, and substantially altered starch particle morphology, crystal structure, tuber yield, and plant growth characteristics (Zhao *et al*., 2021). Harris and Warren utilized Desiree as a model plant and employed Agrobacterium-mediated CRISPR/Cas9 technology to generate various SBE mutants, demonstrating that SBEI and SBEII genes regulate starch branching degree, gelatinization properties, retrogradation characteristics, and starch digestibility (Harris and Warren, 2024).

The tuber starch accumulation genetic loci were also uncovered. Li *et al*. utilized local tetraploid potato cultivars and German tetraploid potato clones as experimental materials, employing starch metabolism functional genes for linkage mapping and developing the SSCP marker Stp28-8b associated with high tuber starch content (Li *et al*., 2013). Fan Shuhua *et al*. conducted genotyping on potato high amylopectin mutant P24023, approved varieties Kexin 12, Zao Dabai, Dongnong 303, Shepody, Eugene, Atlantic, and the F2 hybrid population of P24023 × Kexin 12 using high-resolution melting curve (HRM) molecular marker technology, successfully developing high amylopectin linkage HRM markers (gbss-F/gbss-R) (Fan *et al*., 2019). Jin Xinghong *et al*. studied tetraploid potato hybrid F2 isolation populations and employed mixed grouping analysis techniques to develop two SSR molecular markers SSR-152 and SSR-174 closely linked to potato starch content (Jin *et al*., 2023). Schönhals *et al*. conducted a GWAS using an 8.3K SolCAP chip on 282 tetraploid potato germplasm samples, identifying association regions for starch content across all 12 chromosomes and confirming a major effect locus on chromosome 5, which yielded diagnostic SNP markers for starch yield improvement (Schönhals *et al*., 2016). Li *et al*. integrated SLAF-seq data with BSA pooled association analysis to localize a candidate starch region spanning 0–5.62 Mb on chromosome 2, annotated 41 nonsynonymous starch-related genes, and developed two robustly validated SNP-CAPS functional markers for starch trait characterization, enabling molecular-assisted screening of high-starch materials during seedling development (Li *et al*., 2021). Khlestkin *et al*. conducted a genome-wide association study (GWAS) using a 22K SNP chip on 90 Russian potato germplasms, systematically identifying for the first time loci associated with starch particle morphology and detecting 53 significantly correlated SNPs distributed across nine chromosomes (1, 2, 4, 5, 7, etc.). Notably, eight tightly linked SNPs on chromosome 1 were found to simultaneously regulate both starch extraction rate and particle roundness (Khlestkin *et al*., 2020). In a parallel study published in 2019, Khlestkin’s team identified 17 significant SNPs related to starch phosphorus content, corresponding to eight distinct genomic regions, confirming that phospho-modification of starch exhibits stable, heritable natural variation (Khlestkin *et al*., 2019). Ahmad *et al*. (2026) conducted a GWAS using 102 multi-environment potato germplasms, combining RVA and DSC assays to measure gelatinization and retrogradation parameters; their results identified AGPL on chromosome 1 as controlling gelatinization onset temperature, GWD on chromosome 5 as regulating gelatinization termination temperature, while AGPS on chromosome 7 and SBE jointly determine starch retrogradation characteristics (Ahmad *et al*., 2026). Carpenter *et al*. performed candidate gene association analyses on 193 potato lines, demonstrating that SNPs in the *GWD, SBEI, SBEII*, and *SSSIII* genes regulate C6-and C3-phosphorylation levels of starch respectively, providing key targets for viscosity improvement (Carpenter *et al*., 2015). Schreiber *et al*. identified functional SNPs in eight key starch-sugar interaction genes—including *PHO1* and invertase invGE—through candidate gene association analysis, demonstrating that favorable alleles can simultaneously increase tuber starch content and reduce reducing sugar accumulation during low-temperature storage (Schreiber *et al*., 2014).

Compared to the molecular regulatory networks of starch accumulation in *Arabidopsis* and rice, the molecular regulatory network of starch in potato remains less thoroughly elucidated. Nevertheless, it still needs to be refined through multi-omics data analysis to complete the mining of starch regulatory genes. Key genes were utilized to elucidate starch formation mechanisms and genetically improve commercial varieties, achieving rapid breeding of high-starch potato cultivars.

## 2 Materials and methods

### 2.1 Materials

The experimental materials were hybrid offspring produced by crossbreeding potato cultivars Huasheng No.7, as the female parent, with eight major domestic cultivars – Dongnong 311, Chuntu 11, Kexin 13, Yunshu 306, Yunshu 506, Longshu 7, Kexin 32, and Chuntu 12 – as male parents(Fig. S1). The hybrid progeny included 14 materials with outstanding agronomy traits were selected to identify the starch content in tubers. The experimental materials were growth in Hulunbuir experimental field, which was located at Fuxing Village, Zhonghe Town, Zhalantun City, Inner Mongolia Autonomous Region (48°0′ N, 122°44′ E). The experimental site features hilly terrain with an elevation of 306.5 meters and a calcium-rich loam soil type. The previous crop was corn, and land preparation involved autumn plowing, harrowing, and ridge formation. Field management commenced on April 30,2025, with manual sowing followed by mechanical soil covering and compaction; harvest occurred on September 11, 2025. Fertilization combined basal and topdressing applications: 45 kg/acre of compound fertilizer (12:18:15) plus 10 kg/acre of potassium sulfate as basal fertilizer, supplemented with 25 kg/acre of compound fertilizer (14:5:27) on May 27. Cultivation and weed control measures included deep loosening on May 5, intertillage on May 27, application of herbicide spray on May 29, deep loosening again on June 9, intertillage on June 28, weed removal on July 13, and final weed control on July 30. Irrigation was performed via sprinkler system on May 19, with no manual drainage measures implemented. The harvest tubers were stored at 10°C for 6 months before sample collection, the tubers remain healty and be without significant damage.

### 2.2 Starch content identified

Tuber starch content was estimated via the hydrostatic weighing method (Reiman–Parow method) (Haase, 2003). Undamaged tubers with uniform size (40–60 mm) were cleaned and surface-dried. Each tuber was weighed in air to record W_air_. Subsequently, the tuber was fully suspended in distilled water without touching the container, and the underwater apparent weight W_water_ was recorded. Tuber specific gravity (SG) was calculated as: SG=(W_air_-W_water_)/W_air_. Fresh starch percentage was calculated using the standard empirical regression equation for potato tubers: Starch (% FW)=210.79×SG−213.93. Five tubers were measured per biological replicate, with three replicates set for each treatment.

### 2.3 Sampling collection

Sampling was conducted at tuber bulking, maturation, and storage stages. At each stage, 12 plants were ramdomly selected, and each 3 plants were pooled to one biological replicate. Totally four biological replicates were collected per material. A 0.25–0.5 mm² area on the transverse section of the tuber was selected, encompassing both phloem and xylem tissues. A single-use scalpel blade was used to cut 1–1.5 mm thick sheet samples, which were promptly preserved in 1.5 ml EP tubes using RNA later™ stabilizer (Thermo Fisher Scientific Inc., AM7021) (Fig. S2). The samples were collectively stored at –80°C. During transportation, sufficient dry ice was ensured in the container to avoid RNA degradation.

### 2.4 RNA-Seq Data analysis

Transcriptome sequencing was performed by Novogene Biotech Co., Ltd. using the Illumina sequencing platform. Raw data were removed reads containing adapters, eliminated reads with uncertain bases (N), and deleted low-quality reads (reads with base counts ≤5, Q values exceeding 50% of read length). The filtered data (clean data) undergoes calculation of Q20, Q30 quality scores, error rates of 1% and 0.1%, respectively. The potato genome exhibited a GC content of 34.8%. The index was constructed and reads were aligned to reference genome using HISAT2 v2.2.1 software.The featureCounts v2.0.6 was used to count reads for each gene and calculate FPKM values to quantify gene expression levels. Differential expression analysis was performed using DESeq2 v1.42.0, differentially expressed genes were identified based on adjusted P-values ≤0.05. GO and KEGG enrichment analyses were performed using clusterProfiler v4.8.1 software to identify significantly enriched biological processes and pathways.

### 2.5 Real-time quantitative polymerase chain reaction (qRT-PCR) validate genes expression

The RNA reverse transcription to cDNA procedure was performed according to the instructions of the Evo M-MLV RT Mix Kit with gDNA Clean for qPCR Ver.2 kit. All operations were conducted under ice conditions. A 20μL reverse transcription reaction mixture was prepared, with the maximum RNA usage amount specified in the kit instructions. The reaction protocol included 15-minute incubation at 37°C followed by 5-second denaturation at 85°C. After reaction completion, the mixture was cooled on ice. The resulting cDNA products were stored short-term at-20°C.

The total reaction volume for real-time quantitative PCR (qPCR) detection was 20 μL, with the specific composition ratio as follows: 2× SYBR qPCR Master Mix 10 μL, upstream primers (10 µM) 0.4 μL, downstream primers (10 µM) 0.4 μL, and cDNA template 1 μL. The volume was adjusted to 20 μL with RNase Free Water. The reaction was performed on a fluorescence quantitative PCR instrument under the following conditions: 95°C pre-denaturation for 30s, followed by 1 cycle; 95°C denaturation for 5s, 60°C annealing/extension for 30s, repeated 40 times; 95°C denaturation for 15s, 60°C annealing for 60s, and 95°C amplification at 0.05°C/s per cycle. Relative gene expression was calculated using the Fold Change = 2^-Ct^ method.The amplification primers are listed in Table S1.

### 2.6 Resequencing of Yunshu 306 gDNA and Its Hybrid Offspring

The potato leaf sample DNA was extracted by CTAB method, fellow the publicated protocol (Doyle and Doyle,1987), The genomic DNA resequencing in this study was completed by Annoroad Gene Technology (Beijing) Co., Ltd. utilizing the MGI sequencing platform. FastQC software was employed to perform quality assessment on the raw FASTQ sequencing data. Trimmomatic software was used to clean the raw sequencing reads.The BWA-MEM algorithm was employed to align Clean Reads against the species reference genome. The GATK software is employed to perform base quality recalibration (BaseRecalibration) on the preprocessed BAM files, completing the base quality correction process. Whole-genome SNP variant detection utilizes the GATK HaplotypeCaller algorithm to identify genome-wide SNVs and InDel variants in samples, determine their genotypes, and generate raw VCF variant files.The original variant sites were screened using the following criteria: QUAL ≥ 30 and DP ≥ 10. The population samples underwent re-calculation of variant results using the GATK VQSR model.The ANNOVAR software was employed to perform genomic functional annotation on the filtered high-quality SNP sites.

### 2.7 Detection of β-glucosidase activity

The β-glucosidase (β-GC) activity was determined using a β-glucosidase activity detection kit (Solarbio, product code: BC2560), with samples collected and absorbance measured according to the instructions. First prepare the standard working solution: take 100 μL of 5 μmol/mL p-nitrophenol standard stock solution and add it to 400 μL of distilled water to prepare a 1000 nmol/mL standard solution; then take 100 μL of this solution and add it to 900 μL of distilled water to obtain a 100 nmol/mL standard solution. Weigh approximately 0.2 g of potato tuber sample, add 1 mL of extract solution, fully grind the mixture on ice, centrifuge at 15000 g and 4 for 20 min, and retain the supernatant for testing on ice. Preheat the spectrophotometer for more than 30 min, set the wavelength to 400 nm and calibrate zero with distilled water. Add 500 μL Reagent and 100 μL potato tuber supernatant to the control tube; add 400 μL Reagent, 500 μL Reagent and 100 μL potato tuber supernatant to the test tube. Mix thoroughly and incubate at a constant temperature of 37 for 30 min, then immediately place the tubes in a boiling water bath for 5 min (seal the tubes to avoid liquid splashing). Cool the tubes under running water and mix evenly, then supplement each tube with 400 μL Reagent, centrifuge at 8000 g and 4 for 5 min and collect the supernatant. Next, add 500 μL of the corresponding supernatant to the control tube and test tube respectively, add 500 μL of 100 nmol/mL standard solution to the standard tube, and add 500 μL of distilled water to the blank tube. Add 1000 μL Reagent to all tubes, mix well and leave at room temperature for 2 min, then measure the absorbance of each tube at 400 nm.The calculation formula for enzyme activity is as follows:β-GC activity 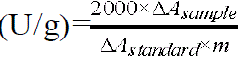, where ΔA_sample_ is the absorbance difference between the sample tube and the control tube, ΔAstandard is the absorbance difference between the standard tube and the blank tube, and m represents the fresh weight of the sample.

## 3 Results and Analysis

### 3.1 Starch concentration in the tuber

The offspring of the Huasheng No7 and other domestical varieties were with starch content ranging from 11.32% to 18.93%.According to their starch content, ten materials were selected, which have the terminal starch concentration in the tuber among the population. Their starch contents were 18.93%, 17.15%, 17.09%, 16.17%, 15.67%, 11.33%, 11.35%, 11.45%, 11.93%, and 12.33% respectively. The ten material groups were numbered as HS1, HS2, HS3, HS4, HS5 and LS1, LS2, LS3, LS4, LS5 respectively. The sampling name in tuber bulking, maturation and storage were shown the table 1.

**Table 1.**
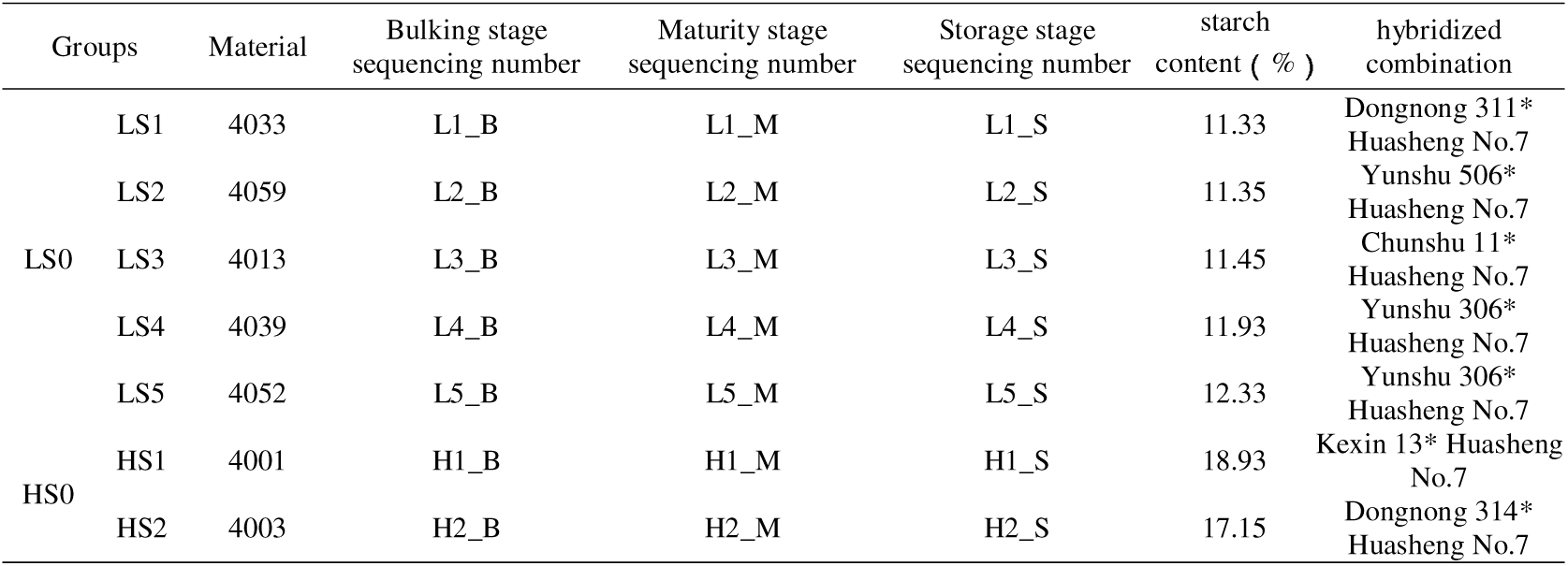
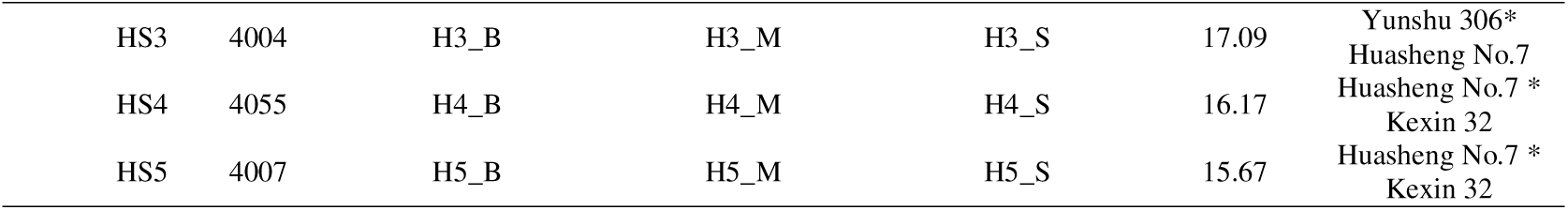
Starch content of 10 hybrid progenies derived from Huasheng No.7 crossed with different potato varieties.

### 3.2 Transcriptome Sequencing Results

The samples generated an average of 46.38 M raw data, with 45.02 M effective data obtained after filtering (Table S3). The Q20 quality scores were consistently above 99%, and the Q30 quality scores were consistently above 97%, indicating reliable sequencing results suitable for subsequent analysis. The average GC content of sequencing samples was 43.57%. Compared to the reported GC content of 34.8% in the potato genome, this high value may reflect the preferential expression of GC-enriched genes in transcriptionally active regions. Correlation heatmaps of transcriptome data for potato tuber samples at three developmental periods (tuber bulking, maturation, storage) were shown in Fig. S4, in which biological replicates within each period displayed correlation coefficients close to 1 with prominent red blocks, while inter-group samples presented relatively lower correlation coefficients with predominantly blue blocks, verifying reliable sample reproducibility and distinct transcriptional differences between treatments. The boxplots of gene expression levels in potato tuber samples were plotted during the tuber bulking stage (Fig. S5A), maturation stage (Fig. S5B), and storage stage (Fig. S5C) to compare and analyze expression differences and their distribution characteristics among different samples. The analysis results indicated that there were no significant deviations in gene expression levels across all samples, suggesting stability in gene expression during these three growth stages.

To elucidate the overall differences in gene expression and biological reproducibility among potato tuber samples at different stages, principal component analysis (PCA) was conducted on samples collected during the bulking stage, maturation stage, and storage stage (Fig. 1). The results demonstrated that intra-group biological replicates across all three stages exhibited tight clustering with no significant outliers, indicating robust experimental reproducibility and high data reliability. Samples from the bulking stage showed relatively concentrated distribution with minimal transcriptomic differences between varieties; certain varieties were closely clustered in PCA space. The H5 group exhibited excessive dispersion, H5 samples from tuber bulking stages were not included in the fellow analysis. Samples from the maturation and storage stages displayed more dispersed distributions, with distinct and non-overlapping clusters along principal component dimensions, reflecting significant inter-varietal gene expression differences. Principal components PC1 and PC2 accounted for 19.13% and 9.26% of gene expression variability respectively at the bulking stage, 14.06% and 13.01% at maturation stage, and 19.63% and 11.68% at storage, both effectively capturing major transcriptional variations across time points. These findings indicate substantial differences in transcriptomic profiles among potato materials at different developmental and storage stage, with materials-specific patterns being particularly pronounced during maturation and storage. The experimental sample grouping was well-designed with reliable data quality, making it suitable for subsequent differential gene identification and functional enrichment analyses.

**Fig. 1.**
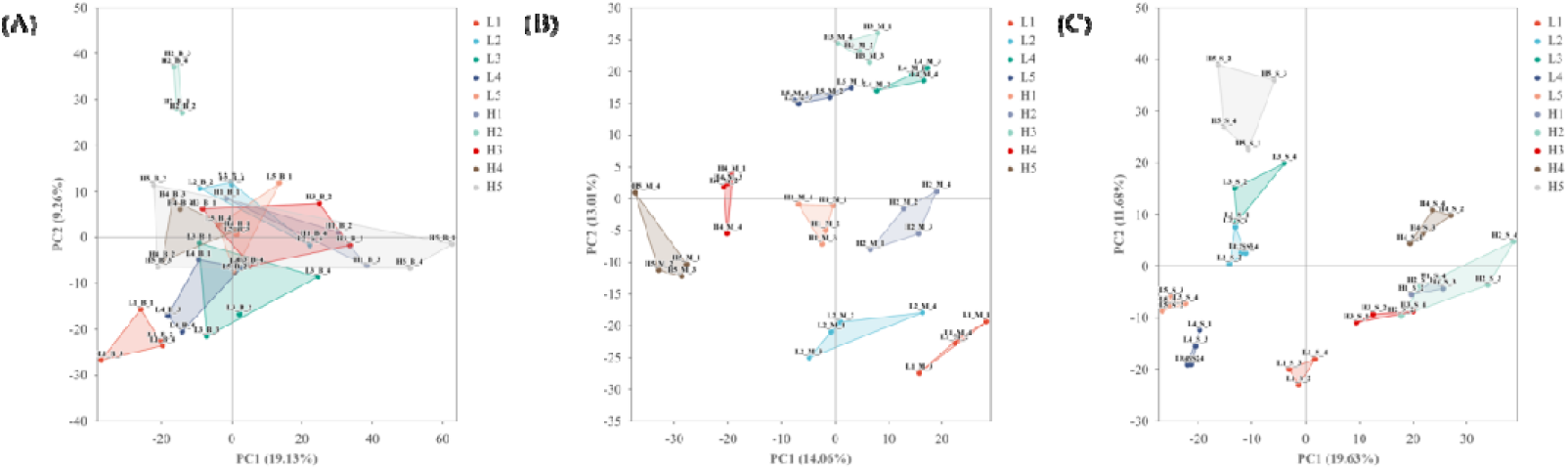
PCA analysis plot of samples during the tuber bulking stage (A), maturation stage(B), and storage stage(C) of potato tubers.

### 3.3 The storage stage had a doubling of differential genes compare to tuber bulking and maturation stage

Transcriptome analysis revealed obvious stage-specific differences in the number of differentially expressed genes (DEGs) between high– and low-starch potato varieties,with low-starch varieties serving as the control group. A total of 559 DEGs (312 upregulated and 247 downregulated; Fig. 2A) were identified at the tuber bulking stage, and 530 DEGs (323 upregulated and 207 downregulated; Fig. 2B) were detected at the maturation stage. The relatively low and stable DEG numbers at these two growth stages indicated that the starch biosynthesis pathway remained stable during potato growth, and continuous starch accumulation did not require extensive genetic reprogramming.

**Fig. 2.**
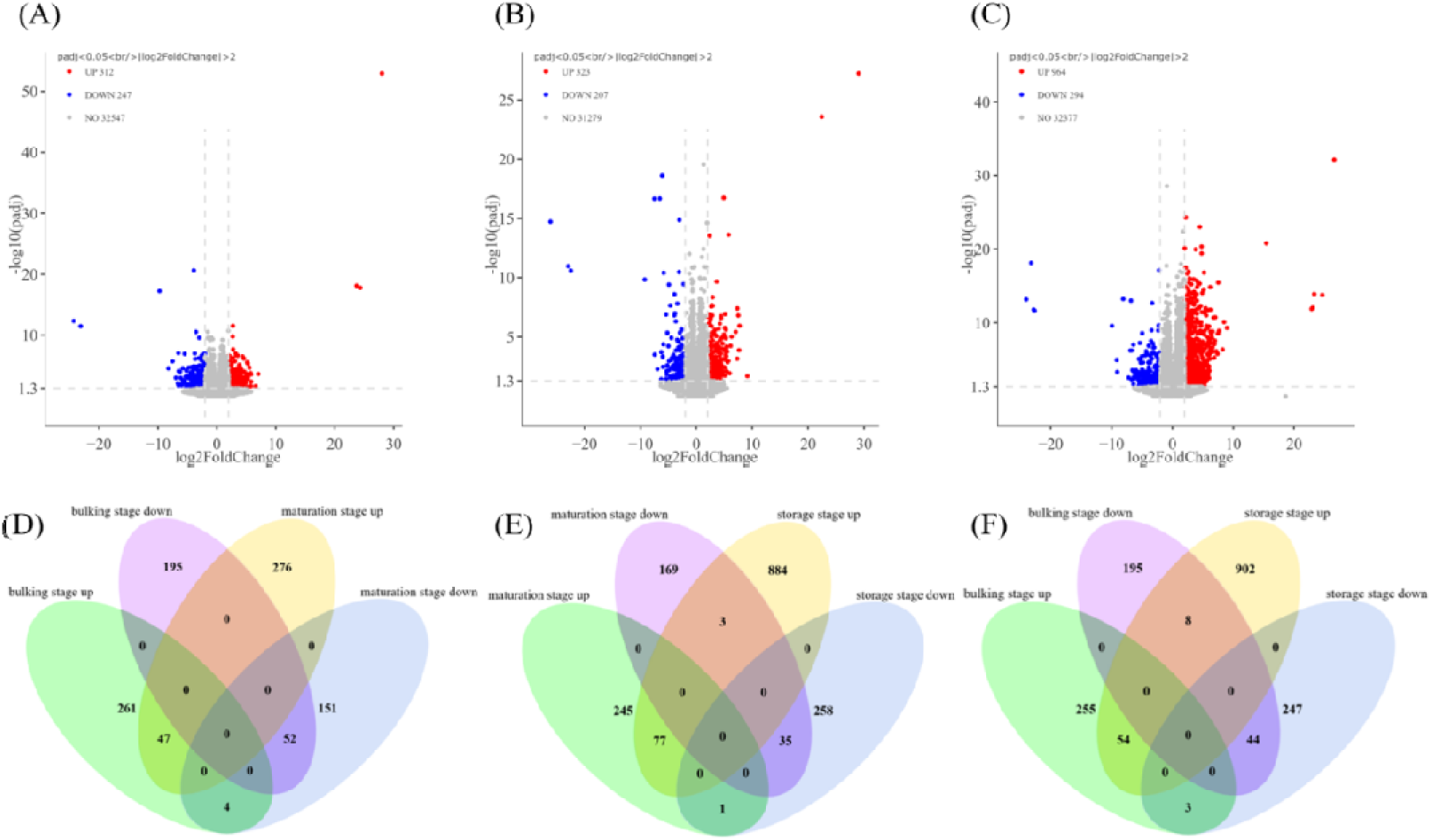
Identification and venn analysis of DEGs in potato tubers at three stages. Note:(A–C) Volcano plots for DEGs (padj<0.05, log2FC >2). Red: up-regulated, blue: down-regulated, gray: non-DEGs. (A) bulking: 312 up, 247 down; (B) maturation: 323 up, 207 down; (C) storage: 964 up, 294 down.(D–F) Venn diagrams of pairwise DEGs. (D) bulking vs maturation; (E) maturation vs storage; (F) bulking vs storage.

In contrast, the number of DEGs increased dramatically to 1,258 (964 upregulated and 294 downregulated; Fig. 2C) at the storage stage, nearly double that of the growth stages. This suggests that numerous physiological defense pathways are activated to adapt to long-term storage conditions. The bulking stage is dominated by cell division and initial starch synthesis with intensive metabolic activity; the maturation stage features peak starch accumulation, which requires the regulation of starch synthesis termination and nutrient allocation; the storage stage relies on genetic regulation to resist environmental stresses, maintain starch quality, and prevent excessive starch degradation (Shirani-Bidabadi *et al*., 2024; Xu and Lian, 1997; Jiang *et al*., 2025; Kumar *et al*., 2024).

Between tuber bulking and maturation stages(Fig. 2D), 99 DEGs are exhibited in both stages, including 47 up-regulated and 52 down-regulated co-expressed genes. 261 genes showed up-regulation and 195 genes down-regulation during tuber bulking specifically. At maturation stage, 276 genes were up-regulated while 195 genes were down-regulated specifically. 4 genes were up-regulated during tuber bulking but down-regulated at maturation. The 456 DEGs were specific found at tuber bulking stage, accounted for 81.57% of total differential genes in this stage, whereas 427 DEGs with specific expression at maturation stage, accounted for 80.56% of total differential genes at this stage, indicated that the starch metabolism pathway different at tuber bulking and maturation stage.

There are 112 some DEGs, including 77 up-regulated and 35 down-regulated genes at the harves and storage stages (Fig. 2E). And 245 genes showed up-regulation, with 169 down-regulation at maturation stage specifically, while 884 DEGs exhibited up-regulation with 258 down-regulation at storage stage. There is only one genes were up-regulated during at maturation but down-regulated at storage, and 3 genes were down-regulated at maturation but up-regulated at storage. At maturation stage, 408 DEGs were found specifically, accounting for 76.98% of total different expression genes at this stage, whereas at storage stage, 1,142 specifically found, representing 90.78% of at this stage. This results indicate that there are some same type DEGs at tuber maturation and storage stages, but most are the different DEGs, the starch metabolism share some similar pathway, but more especially.

There are 98 some DEGs were identified, including 54 upregulated and 44 downregulated genes at tuber bulking and storage stages (Fig. 2F). Specifically, 255 DEGs exhibited upregulation during tuber buling,while 195 genes showed downregulation. At storage stage, 902 genes were upregulated and 247 genes were downregulated, especially. Three genes were upregulated during tuber bulking but downregulated during storage, while eight genes were downregulated during tuber bulking but upregulated during storage. Totally, 450 genes were specifically expressed at tuber bulking stage, accounting for 80.5% of all DEGs at this stage, whereas 1,149 DEGs were specifically at storage stage, representing 91.33% of the entire DEGs.

### 3.4 Functional enrichment analysis of genes differentially expressed during potato tuber development and storage mediated by cell structure and carbohydrate metabolism

Gene Ontology (GO) enrichment analysis was performed on DEGs isolated from tuber samples at bulking, maturation and storage stages (Fig. 3). Distinct enrichment patterns across developmental phases were observed. DEGs at the tuber bulking stage were significantly enriched in 5 cellular component (CC) terms and 8 molecular function (MF) terms; DEGs from the maturation stage were enriched in 5 CC terms and 5 MF terms; DEGs from the storage stage exhibited significant enrichment in 15 biological process (BP) terms, 5 CC terms and 6 MF terms.

**Fig. 3.**
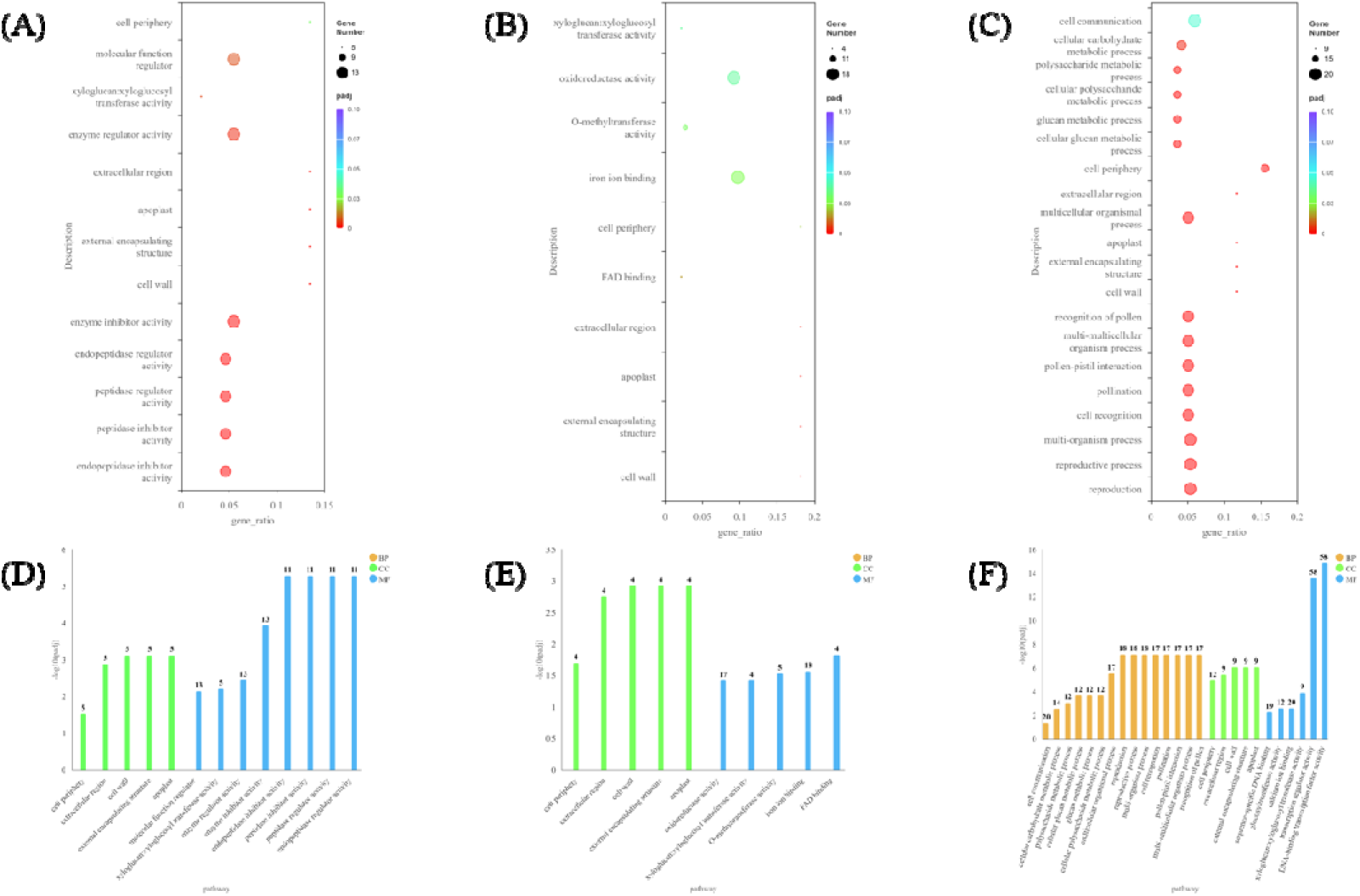
GO Enrichment Results of Differentially Expressed Genes in Potato Tubers at Three Developmental Stages. Note: a, b and c represent the bubble plots of GO enrichment at the tuber bulking stage, maturation stage and storage stage, respectively; d, e and f represent the bar charts of GO enrichment at the tuber bulking stage, maturation stage and storage stage, respectively.

Across all three stages, five consistent CC terms were repeatedly enriched, including cell wall, external encapsulating structure, apoplast, extracellular region and cell periphery. Xyloglucan:xyloglucosyl transferase (XET) activity was the core shared MF term among all stages. Genes annotated to XET activity encode proteins that mediate cleavage and reconnection of xyloglucan chains, which regulates cell wall relaxation, cell elongation and cell wall structural reconstruction (Xiong *et al*., 2026; Yuan *et al*., 2024; Seale, 2020).

#### 3.4.1 Tuber bulking stage

The five universal CC terms shared a unified set of five DEGs at the bulking stage: three upregulated transcripts (*Soltu.DM.07G022400,Soltu.DM.03G015050,Soltu.DM.07G021990*) and two downregulated transcripts (*Soltu.DM.02G027190, Soltu.DM.07G005220*). These five DEGs were also assigned to the XET activity MF term. XET proteins encoded by these genes modulate the spatial structure of peripheral tissues and facilitate the translocation and partitioning of photosynthates to sink tubers (Hidvégi *et al*., 2024). No significantly enriched BP terms were identified at this stage.

#### 3.4.2 Tuber maturation stage

Four identical upregulated DEGs were detected within all five CC terms at maturation:*Soltu.DM.12G028730, Soltu.DM.07G021990, Soltu.DM.10G000430, Soltu.DM.12G028740*. These four genes were annotated to XET activity. Continuous expression of these XET coding genes maintains intact cell wall architecture and alleviates oxidative damage to intracellular metabolic enzymes (Shirani-Bidabadi *et al*., 2024). No significantly enriched BP terms were found at the maturation stage.

#### 3.4.3 storage stage

Nine uniformly upregulated DEGs co-existed in four CC terms (cell wall, external encapsulating structure, apoplast, extracellular region): *Soltu.DM.03G015050*, *Soltu.DM.03G015070, Soltu.DM.12G028730, Soltu.DM.12G028740, Soltu.DM.03G015060, Soltu.DM.03G015040, Soltu.DM.12G020770, Soltu.DM.07G022400, Soltu.DM.10G000430*. These nine transcripts were also enriched in the XET activity MF term. Sustained XET-mediated cell wall remodeling coordinates carbohydrate turnover, disease defense and vitamin B6 metabolism to stabilize cellular structure (Wang *et al*., 2026; Zhou *et al*., 2020).

When comparing high-starch and low-starch tuber materials, a total of 12 upregulated genes were specifically enriched in the cell periphery CC term: *Soltu.DM.06G031080, Soltu.DM.03G015050, Soltu.DM.03G015070, Soltu.DM.12G028730, Soltu.DM.12G028740, Soltu.DM.03G015060, Soltu.DM.03G015040, Soltu.DM.12G020770, Soltu.DM.09G007290, Soltu.DM.09G007300, Soltu.DM.07G022400, Soltu.DM.10G000430*.

Different from bulking and maturation tubers with no enriched BP entries, five carbohydrate-related BP terms were significantly enriched by upregulated DEGs in storage tubers, including cellular glucan metabolic process, glucan metabolic process, cellular polysaccharide metabolic process, polysaccharide metabolic process, and cellular carbohydrate metabolic process. Twelve consistently upregulated genes responsible for cell wall polysaccharide synthesis and remodeling were screened out, including three cellulose synthase-like (CSL) genes (*Soltu.DM.03G020780, Soltu.DM.07G014150, Soltu.DM.08G003680*) and nine XET/XTH family genes. Their protein products reconstruct cell wall polysaccharide networks and reshape the pool of intracellular soluble carbohydrates (Sergeeva *et al*., 2022; Zhang *et al*., 2017).

Across bulking, maturation and storage stages, five DEGs displayed persistent differential expression throughout the whole developmental and storage cycle: *Soltu.DM.07G022400, Soltu.DM.03G015050, Soltu.DM.12G028730, Soltu.DM.12G028740, Soltu.DM.10G000430*. Among them, *Soltu.DM.07G022400* and *Soltu.DM.03G015050* showed differential expression only at the bulking and storage stages, while the remaining three genes maintained stable upregulation from maturation to storage.

### 3.5 Analysis of starch homeostasis mechanisms during the developmental and storage stages of potato tubers mediated by multidirectional metabolic regulation

#### 3.5.1 Tuber Bulking Stage

KEGG pathway enrichment analysis revealed that differentially expressed genes (DEGs) at the tuber bulking stage were significantly enriched in three pathways: photosynthesis–antenna proteins (Fig. S6a), glutathione metabolism (Fig. S6c), and plant circadian rhythm (Fig. S6b). The photosynthesis–antenna protein pathway contained seven upregulated DEGs, including five genes encoding chlorophyll a/b-binding proteins (*Soltu.DM.02G014010, Soltu.DM.02G013810, Soltu.DM.03G000850, Soltu.DM.03G000840, Soltu.DM.03G000860*), one gene encoding photosystem I biogenesis subunit A4 (*Soltu.DM.06G025820*), and one gene encoding photosystem I assembly protein (*Soltu.DM.12G028930*). This canonical pathway functions to capture light energy and sustain plant photosynthetic carbon fixation (Shi *et al*., 2024).

A total of nine DEGs were enriched in the glutathione metabolism pathway, including eight glutathione S-transferase family genes (*Soltu.DM.10G023750, Soltu.DM.09G001280, Soltu.DM.12G017800, Soltu.DM.01G028030, Soltu.DM.09G001400, Soltu.DM.09G001320, Soltu.DM.09G001360, Soltu.DM.12G028950*) and one gene encoding cytoplasmic aminopeptidase (*Soltu.DM.12G027320*), among which seven were upregulated and two were downregulated. Glutathione metabolism is a core antioxidant pathway that eliminates excess intracellular ROS and maintains cellular redox homeostasis (Naher *et al*., 2024).

Five DEGs were enriched in the plant circadian rhythm pathway, including two upregulated DEGs encoding GIGANTEA (*Soltu.DM.04G027760*) and cryptochrome synthase family protein (*Soltu.DM.09G028560*), and three downregulated DEGs encoding pseudo-response regulators (*Soltu.DM.10G012570, Soltu.DM.11G012010*) and early-flowering protein ELF3 (*Soltu.DM.06G022810*). This pathway mediates endogenous circadian clock oscillation and regulates the temporal expression of diverse plant metabolic genes to coordinate plant growth and environmental adaptation (Wu *et al*., 2015).

#### 3.5.2 Tuber Maturation Stage

At the tuber maturation stage, DEGs were significantly enriched in four pathways: phenylpropanoid biosynthesis (Fig. S7a), fatty acid elongation (Fig. S7b), photosynthesis–antenna proteins (Fig. S7d), and fatty acid biosynthesis (Fig. S7c). The phenylpropanoid biosynthesis pathway included seven upregulated and three downregulated DEGs, which primarily encoded phenylalanine ammonia-lyase (*Soltu.DM.09G005710, Soltu.DM.09G005700, Soltu.DM.09G005690, Soltu.DM.03G011440*) and peroxidase proteins (*Soltu.DM.09G009900, Soltu.DM.05G018820, Soltu.DM.01G001850*), and it participates in the synthesis of lignin and phenolic secondary metabolites and responds to cellular oxidative damage (Navarre *et al*., 2013). Fatty acid elongation and fatty acid biosynthesis pathways shared five uniformly upregulated DEGs encoding acyl reductases (*Soltu.DM.12G003600, Soltu.DM.12G003620, Soltu.DM.12G003560, Soltu.DM.12G003640, Soltu.DM.12G003580*), which are responsible for the synthesis and extension of cellular membrane lipids (Zhang *et al*., 2023). In addition, four upregulated DEGs encoding chlorophyll a/b-binding proteins (*Soltu.DM.02G014000, Soltu.DM.12G023640, Soltu.DM.03G030020, Soltu.DM.02G014050*) were enriched in the photosynthesis–antenna protein pathway during tuber maturation, and their expression dynamics are closely associated with plant photosynthetic performance during tissue senescence (Geigenberger *et al*., 2004).

#### 3.5.3 Storage Stage

During storage stage, DEGs enriched in the plant–pathogen interaction pathway (Fig. S8a) encoded nucleotide-gated channel (*Soltu.DM.05G020050*), calmodulin-like proteins (*Soltu.DM.10G018290, Soltu.DM.02G007250, Soltu.DM.11G024600, Soltu.DM.11G024620, Soltu.DM.11G024610, Soltu.DM.02G034080, Soltu.DM.06G029230*), and EF-hand calcium-binding proteins (*Soltu.DM.02G031620, Soltu.DM.10G027990, Soltu.DM.01G019360, Soltu.DM.02G026800, Soltu.DM.03G033180, Soltu.DM.01G007760, Soltu.DM.04G024930*), which collectively form a complete calcium signal perception and transduction system. A total of 22 DEGs were upregulated and one gene (*Soltu.DM.12G028030*) was downregulated in this pathway, including three WRKY transcription factor genes: StWRKY8 (*Soltu.DM.06G018840*), StWRKY9 (*Soltu.DM.09G009490*), and StWRKY22 (*Soltu.DM.10G005890*). This pathway regulates multi-layered stress responses involving calcium signaling, ROS signaling, and membrane immune recognition (Khan *et al*., 2026). DEGs enriched in the vitamin B6 metabolism pathway (Fig. S8b) were five upregulated phosphate starvation-responsive genes (*Soltu.DM.03G036180, Soltu.DM.03G036190, Soltu.DM.03G036170, Soltu.DM.06G022750, Soltu.DM.06G022730*), which regulate intracellular phosphate transport and redistribution to ensure the activation of vitamin B6 under low-temperature conditions (Nasr *et al*., 2025).

### 3.6 The DEGs relative to the Starch and Sucrose Metabolic Pathways

To uncover the starch accumulation relative candidate genes, totally 150 genes, which were the homologue genes in potato relative to starch systhesis and degradation from other model plants were identified in potato, which are listed in Table S3. Expression levels of the genes identified in the starch and sucrose metabolism pathway and DEGs from three stages were shown in Fig. 5. Potato tubers undergo dramatic temporal metabolic differentiation across three key developmental phases: swelling, maturation and postharvest storage. A total of 24 genes involved in sugar turnover, cell wall remodeling and starch anabolism/catabolism exhibit strict stage-specific expression profiles, which coordinate to orchestrate tuber morphogenesis, carbon partitioning, starch deposition and final storage-processing quality.

**Fig. 4.**
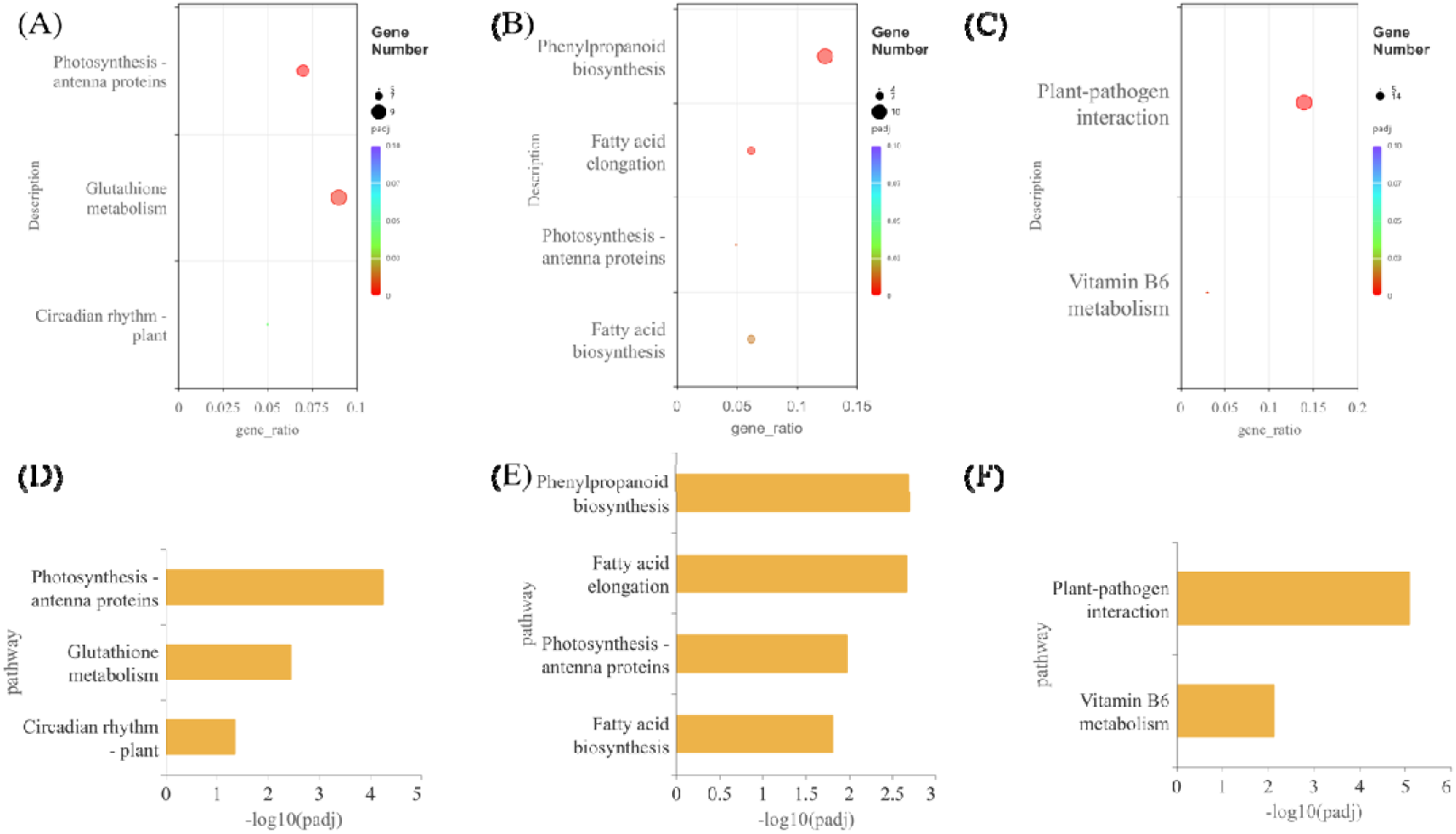
KEGG Enrichment Results of Differentially Expressed Genes in Potato Tubers at Three Developmental Stages. Note: a, b and c represent the KEGG enrichment bubble plots of the tuber bulking stage, maturation stage and storage stage, respectively; d, e and f represent the bar charts of KEGG enrichment at the tuber bulking stage, maturation stage and Storage stage, respectively.

**Fig. 5.**
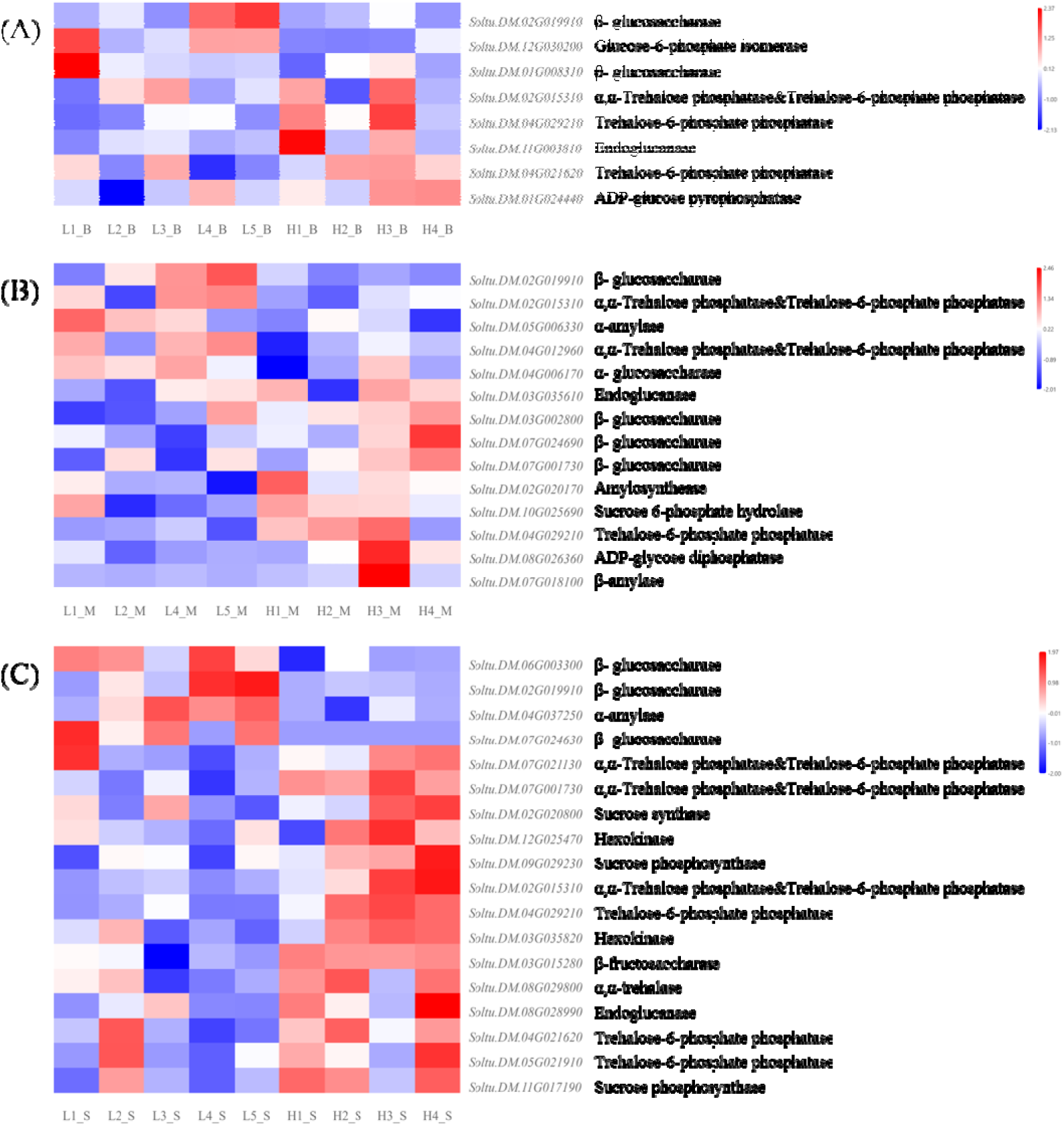
Heatmaps of starch and sucrose metabolic pathway gene expression during potato tuber bulking stage (A), maturation stage (B), and storage stage (C) Enzymes encoded by corresponding genes are shown on the right of each gene.

Genes highly activated during tuber swelling all encode functional enzymes that jointly regulate cell wall loosening, central carbon flux balance and the early supply of starch precursors to sustain rapid cell proliferation and biomass buildup. *Soltu.DM.11G003810* encodes endoglucanase, which degrades cellulosic polysaccharides to relax cell wall rigidity and remove physical barriers to cell expansion and tuber morphogenesis (Nicol *et al*., 1998). *Soltu.DM.12G030200 (GPI)* encodes glucose-6-phosphate isomerase, which catalyzes the reversible interconversion of glucose-6-phosphate (G6P) and fructose-6-phosphate, linking glycolysis, sucrose synthesis and starch biosynthetic pathways to stabilize carbon metabolic flux and ensure sufficient carbon substrates for rapid tuber growth (Gong X *et al*., 2024). *Soltu.DM.01G024440 (agpS3)* encodes ADP-glucose pyrophosphorylase, which maintains stable intracellular pools of ADP-glucose, fine-tunes starch synthesis rates and balances tuber cell growth with early starch deposition (Sowokinos and Preiss, 1982). *Soltu.DM.04G021620* and *Soltu.DM.04G029210* both encode trehalose-6-phosphatase, which modulates trehalose signaling pathways, improves cellular osmotic tolerance and buffers metabolic fluctuations caused by rapid cell division to guarantee orderly metabolism at the swelling stage (Kolbe *et al*., 2005).

Genes predominantly expressed in maturing tubers encode a set of metabolic enzymes that coordinate massive starch biosynthesis, continuous carbon substrate delivery and enhanced intercellular transport to form tuber dry matter and intrinsic textural quality. *Soltu.DM.02G020170 (SS3)* encodes starch synthase, which continuously elongates glucan chains to drive large-scale synthesis of amylose and amylopectin, directly determining tuber dry weight and floury processing traits (Seung *et al.,* 2015). *Soltu.DM.08G026360 (aspP)* encodes ADP-glucose diphosphatase, which modulates starch precursor metabolism and prevents excessive starch synthesis to sustain balanced starch accumulation during maturation (Sowokinos and Preiss, 1982). *Soltu.DM.10G025690 (SPP2)* encodes sucrose-phosphate phosphatase, which catalyzes the terminal step of sucrose biosynthesis, elevating sucrose levels in tubers to provide abundant carbon skeletons for starch production (Stein and Granot, 2019). *Soltu.DM.04G006170* encodes α-glucosidase, which hydrolyzes maltooligosaccharides to release free glucose and adjust soluble sugar profiles. *Soltu.DM.05G006330* encodes α-amylase and *Soltu.DM.07G018100* encodes β-amylase; the two enzymes act synergistically to degrade starch and remodel starch granule structures, balancing tuber sweetness and floury texture (Yang *et al*., 2023). *Soltu.DM.03G035610* encodes endoglucanase, which moderately remodels cell wall structures to facilitate photosynthate translocation. *Soltu.DM.02G019910* encodes β-glucosidase, which fine-tunes cell wall polysaccharide composition, facilitates intercellular solute transport and reshapes intracellular sugar pools (Nicol *et al*., 1998; Cairns and Esen, 2010). *Soltu.DM.04G012960* encodes trehalose phosphate phosphatase and trehalose-6-phosphate hydrolase, which reduce cellular energy consumption and maintain intracellular metabolic homeostasis to support efficient starch synthesis (Kolbe *et al*., 2005). Genes induced during tuber storage encode enzymes that mediate cell wall catabolism, sucrose cycling, clearance of toxic reducing sugars and abiotic stress resistance, controlling tuber softening, senescence and storage quality stability. *Soltu.DM.08G028990* encodes endoglucanase, which continuously degrades cell wall polysaccharides to trigger natural postharvest softening and enable cyclic reuse of cell wall-derived carbon resources (Nicol *et al*., 1998). *Soltu.DM.06G003300* and *Soltu.DM.07G024630* both encode β-glucosidase, which work cooperatively to hydrolyze cell wall glycosides and oligosaccharides and promote reducing sugar accumulation, thus participating in the regulation of postharvest senescence (Cairns and Esen, 2010). *Soltu.DM.03G035820 (HXK1)* encodes hexokinase, which phosphorylates free monosaccharides to eliminate excess harmful reducing sugars, inhibit enzymatic browning and slow quality deterioration during storage (Rodríguez-Saavedra *et al*., 2021). *Soltu.DM.11G017190* encodes sucrose phosphate synthase and *Soltu.DM.02G020800* encodes sucrose synthase; the two enzymes coordinate to maintain endogenous sucrose homeostasis and continuously supply carbon substrates for basal respiratory metabolism (Stein and Granot, 2019). *Soltu.DM.07G001730* and *Soltu.DM.02G015310* encode trehalose metabolic enzymes that mitigate intracellular oxidative damage, stabilize tuber dormancy and boost overall storage stress tolerance (Kolbe *et al*., 2005). *Soltu.DM.04G037250 (AMYA1)* encodes α-amylase, which efficiently degrades aged starch and serves as the core enzyme initiating postharvest soluble sugar remodeling (Yang *et al*., 2023). *Soltu.DM.03G015280 (VInv)* encodes vacuolar β-fructosidase, which specifically hydrolyzes sucrose under vacuolar acidic conditions to produce glucose and fructose. Excess reducing sugars generated by this enzyme induce acrylamide formation during thermal processing, making this gene a critical target for improving potato storage and processing quality (Wiberley-Bradford *et al*., 2014; Zhu *et al*., 2014; Cui *et al*., 2025). Results revealed that two genes, *Soltu.DM.02G019910* and *Soltu.DM.02G015310*, exhibit across all three stages, indicated they may play regulation role in three stages.

There are four module during starch and sucrose metabolic pathway, contained cellulose metabolism, sucrose metabolism, glycoside/trehalose metabolism and starch synthesis. Corresponding pathway maps were constructed based on these candidate genes (Fig. 6). At tuber bulking stage, the cellulose metabolism and glycoside/trehalose metabolism model contain most candidate genes (*Soltu.DM.11G003810,Soltu.DM.02G019910,Soltu.DM.04G029210*), indicated that during the tuber bulking the tuber start to produce more cellulose and trehalose, which be used for tuber bulking, and not for starch synthesis. At maturation stage, all four modules contain the some DEGs, indicated the starch and sucrose metabolic pathways were very activities at this stage, include the starch synthesis and degradation process. Especially, the *Soltu.DM.08G026360*, and *Soltu.DM.02G020170* were upregulated in high starch materials and play role in transfer α-D-Glucose 1P to ADP-glucose, and then to systhesis starch.

**Fig. 6.**
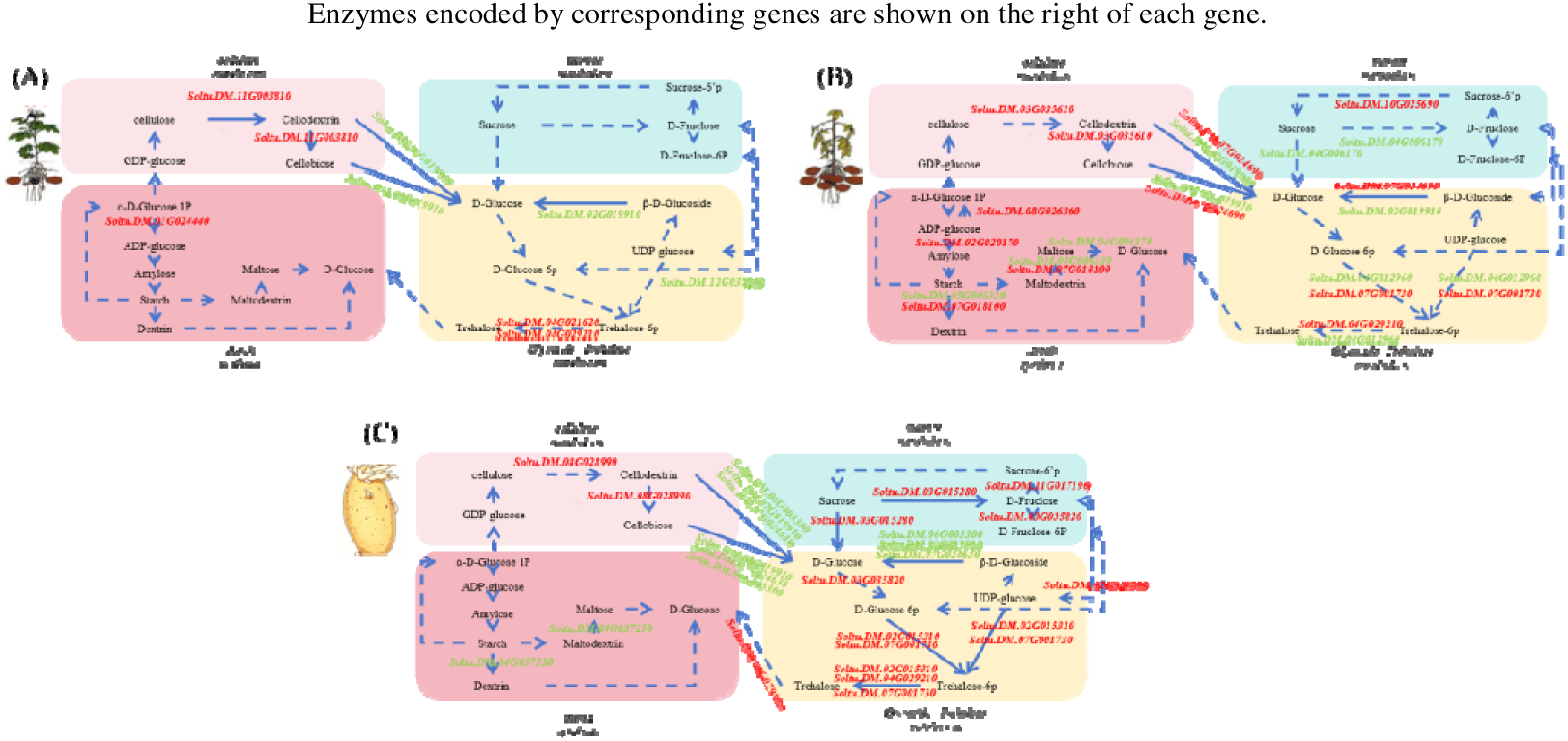
Starch and sucrose metabolic pathways during potato tuber bulking stage (A), maturation stage (B), and storage stage (C)

At tuber storage, most DEGs located in sucrose metabolism, glycoside/trehalose metabolism and cellulose metabolism, produced more D-Glucose 6p, Trehalose-6p and Trehalose via hexokinase ( *Soltu.DM.03G035820*), α,α-Trehalose phosphatase (*Soltu.DM.07G001730, Soltu.DM.02G015310*) and Trehalose-6-phosphate phosphatase (*Soltu.DM.02G015310, Soltu.DM.04G029210, Soltu.DM.07G001730*). Especially, at this stage, very few DEGs located in starch synthesis, which indicated that starch metabolism is relaitve to starch degradation at this stage.

### 3.7 The DEGs between high starch and low starch enrich more SNPs between the parient

DEG was induce by the genenic different between parients.The high-starch materials HS3, low-starch materials LS4 and LS5, along with their parental lines Yunshu 306 and Huasheng No.7 (high starch parent), were subjected to resequencing. Table S4 presents their sequencing quality. All detected SNP loci were further analyzed. Loci with consistent genotypes in the high-starch group (HS3, Huasheng No.7) but divergent genotypes in the low-starch group (LS4, LS5), as well as loci showing the opposite genotypic pattern, were defined as candidate polymorphic SNPs, whereas the remaining non-polymorphic SNPs were excluded. Table 3 was generated based on this screening standard, integrating SNP variation information and differentially expressed genes (DEGs) of starch and sucrose metabolic pathway from potato tuber transcriptomes at three developmental stages.

This result indicate the DEGs on different stage were induced by the SNPs from their parent, and the allele from the high starch parent, Huasheng 7 may contribute the high starch allele to the offsprings, especially for gene *Soltu.DM.03G035610 Soltu.DM.04G029210, Soltu.DM.02G015310, Soltu.DM.02G020800*.

**Table 2.**
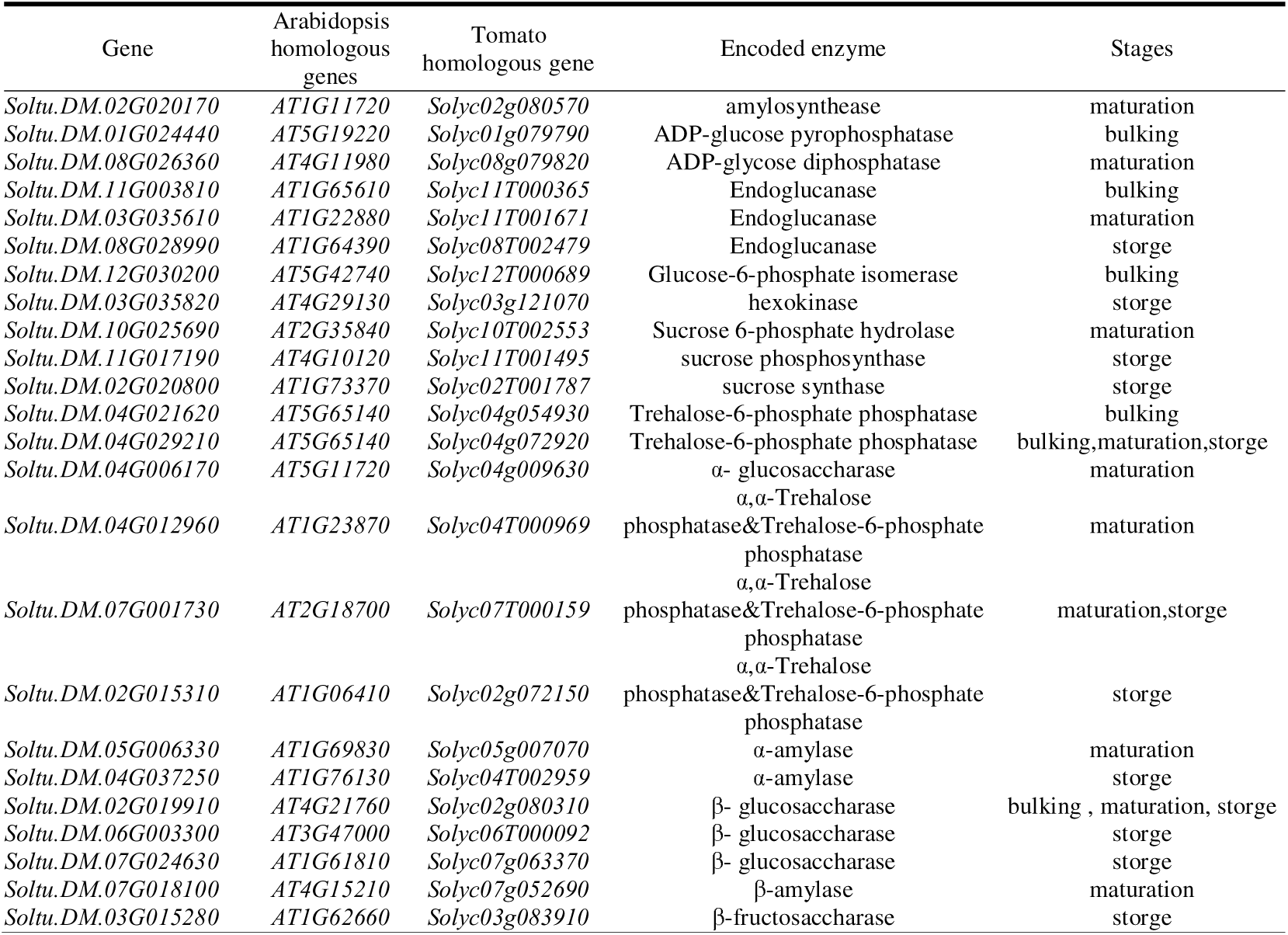
Genes with differential expression in starch and sucrose metabolic pathways across three stages.

**Table 3.** Counts and proportions of polymorphic SNPs in candidate genes related to tuber starch metabolism.

| Gene | Polymorphic SNPs | Total SNPs | Polymorphism |
| --- | --- | --- | --- |
| *Soltu.DM.02G019910* | 20 | 139 | 14.39% |
| *Soltu.DM.05G006330* | 0 | 4 | 0.00% |
| *Soltu.DM.07G018100* | 2 | 24 | 8.33% |
| *Soltu.DM.03G035610* | 13 | 63 | 20.63% |
| *Soltu.DM.04G012960* | 51 | 327 | 15.60% |
| *Soltu.DM.04G006170* | 0 | 20 | 0.00% |
| *Soltu.DM.02G020170* | 7 | 47 | 14.89% |
| *Soltu.DM.04G029210* | 88 | 352 | 25.00% |
| *Soltu.DM.07G001730* | 5 | 90 | 5.56% |
| *Soltu.DM.03G002800* | 2 | 55 | 3.64% |
| *Soltu.DM.11G003810* | 4 | 101 | 3.96% |
| *Soltu.DM.04G021620* | 12 | 579 | 2.07% |
| *Soltu.DM.01G024440* | 1 | 53 | 1.89% |
| *Soltu.DM.01G008310* | 3 | 189 | 1.59% |
| *Soltu.DM.02G015310* | 40 | 127 | 31.50% |
| *Soltu.DM.03G035820* | 17 | 55 | 30.91% |
| *Soltu.DM.03G015280* | 7 | 75 | 9.33% |
| *Soltu.DM.02G020800* | 3 | 9 | 33.33% |
| *Soltu.DM.07G024630* | 0 | 35 | 0.00% |
| *Soltu.DM.04G037250* | 6 | 84 | 7.14% |
| *Soltu.DM.05G021910* | 0 | 40 | 0.00% |

Note: Polymorphic SNPs: Number of filtered candidate polymorphic SNPs ;Total SNPs : Total raw SNP counts detected within each gene from the original datasets ;Polymorphism : Percentage of candidate polymorphic SNPs relative to total raw SNPs in each group.

### 3.8 In low starch group, *Soltu.DM.02G019910* and *Soltu.DM.02G015310* were high and low expression, respectively,with valicated by quantitative real-time PCR (qRT-PCR)

The expression levels of two key genes, *Soltu.DM.02G019910* (encoded β–glucosaccharase) and *Soltu.DM.02G015310* (encoded α,α-Trehalose phosphatase&Trehalose-6-phosphate phosphatase) were detected in samples from three developmental stages via quantitative real-time PCR (qRT-PCR) technology. As shown in Fig. 7, the expression of *Soltu.DM.02G019910* in low starch group at maturation stage (Fig. 7B) and storage stage (Fig. 7C) were significantly higher than high-starch group, and higher in low starch group at tuber bulking stage too (Fig. 7A). In contrast, the expression level of *Soltu.DM.02G015310* was significantly lower in the low-starch group during tuber enlargement (Fig.7D) and storage period (Fig.7F) compared to the high-starch group, with no significant difference at maturity stage (Fig.7E). The expression level results indicated that the β-glucosidase, which *Soltu.DM.02G019910* encoded may have high enzyme activity in the low starch group, and α,α-Trehalose phosphatase&Trehalose-6-phosphate phosphatase enzyme activity in high starch group may higher than low starch group. Meanwhile, these two key genes across all three stages consistently aligned with those derived from transcriptome sequencing, findings conclusively confirm the high reliability and strong reference value of the transcriptome sequencing data in this study, laying a solid experimental foundation for subsequent research based on sequencing data.

**Fig. 7.**
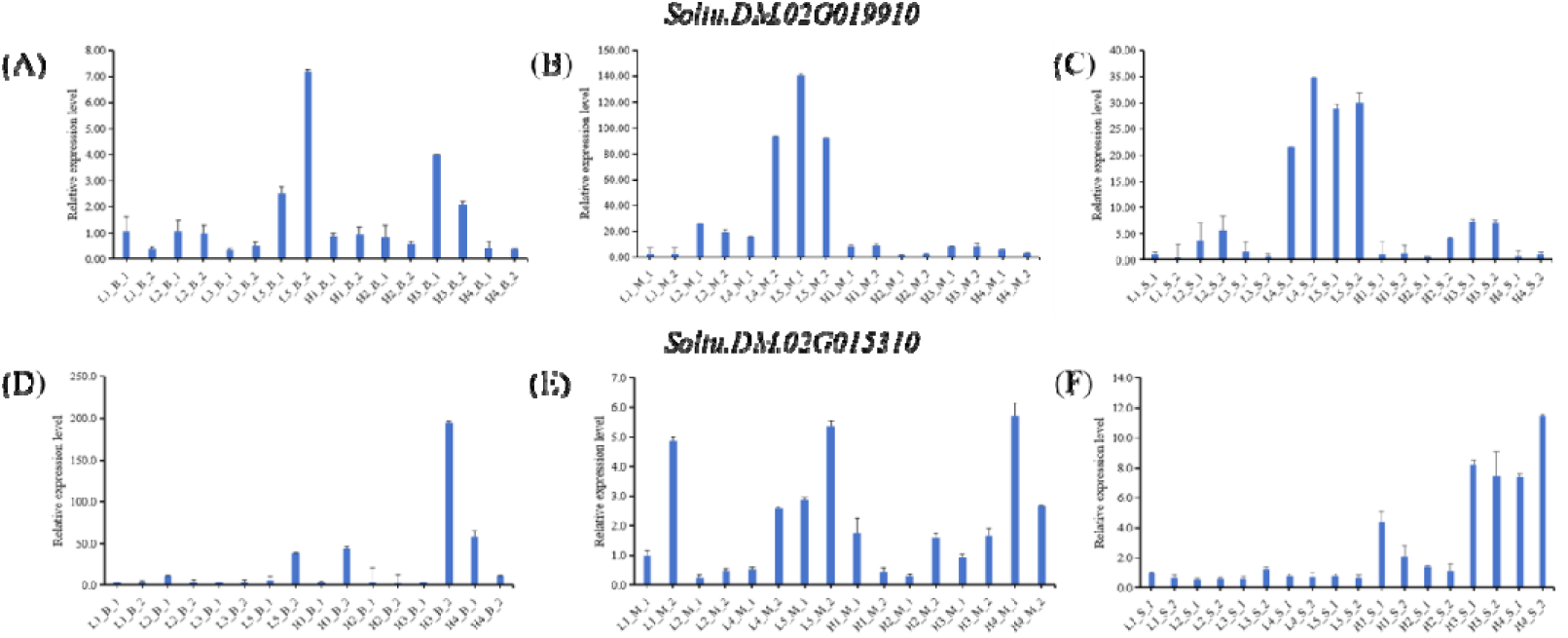
qRT-PCR analysis of *Soltu.DM.02G019910* and *Soltu.DM.02G015310* in potato tubers at three developmental stages a, b and c shown the relative expression levels of *Soltu.DM.02G019910* at the tuber bulking stage, maturation stage and storage stage, respectively; d, e and f shown the relative expression levels of *Soltu.DM.02G015310* at the tuber bulking stage, maturation stage and storage stage, respectively.

### 3.9 In low starch population, the β-glucosidase activity is significantly higher that high starch population

To validate *Soltu.DM.02G019910*’s function, a breeding population’s β-glucosidase Activity was identified. The tuber starch concentration of the population, which contain 24 materials was shown in Table S5, the materials were grouped according to starch content: the high-starch group (HS) with starch content > 12%, and the low-starch group (LS) with starch content < 12%. Each material’ β-glucosidase Activity were identified.The results showed that the overall β-glucosidase activity of the low-starch group was higher than that of the high-starch group significantly (Fig. 8). This pattern of enzyme activity difference concurred with *Soltu.DM.02G019910*’s expression level from previous transcriptome sequencing analyses and RT-PCR result.

**Fig. 8.**
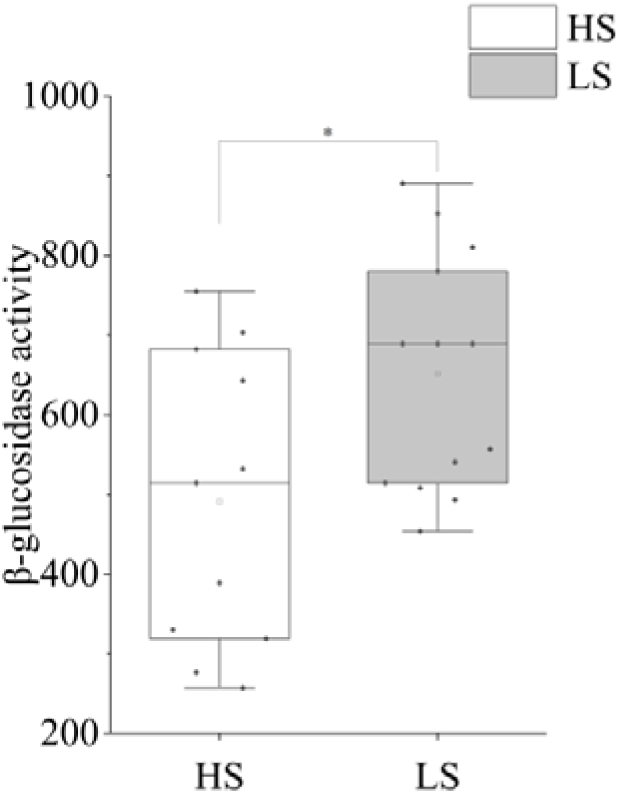
β-glucosidase activity in high and low starch breeding population respecitivlely.

## 4 Discussion

### 4.1 Analysis of Stage-Specific Transcriptional Patterns Underlying Tuber Starch Metabolism

Transcriptome profiling across tuber developmental and storage stages revealed pronounced temporal specificity in global transcriptional reprogramming, which tightly matches the dynamic physiological demands of starch metabolism and environmental adaptation. Overall, tuber growth stages (bulking and maturation) maintained relatively stable and low-level transcriptional variation, whereas postharvest storage triggered drastic transcriptomic alteration, indicating fundamentally distinct regulatory strategies governing starch turnover during growth and storage.

The limited transcriptional variation during tuber bulking and maturation suggests that core starch biosynthetic machinery is relatively conserved and constitutively active during tuber growth. Continuous starch accumulation and tuber morphogenesis do not require large-scale reprogramming of genetic networks, and only a small set of stage-shared genes are sufficient to sustain steady starch anabolism and carbon allocation. Meanwhile, the high proportion of stage-specific DEGs at these two growth phases further demonstrates that tuber bulking and maturation rely on discrete transcriptional modules to drive distinct physiological processes: the former dominates cell proliferation and the initiation of starch synthesis, while the latter controls starch accumulation plateau and growth termination (Shirani-Bidabadi *et al*., 2024; Kumar *et al*., 2024).

In sharp contrast, the substantially elevated transcriptional complexity during storage reflects a comprehensive adaptive response to low-temperature stress and postharvest metabolic remodeling. The massive induction of stage-specific genes indicates that tubers actively activate extensive stress defense, carbohydrate turnover, and metabolic homeostasis pathways to counteract cold-induced starch degradation and quality deterioration (Xu and Lian, 1997; Jiang *et al*., 2025). Although partial genetic overlap exists between maturation and storage stages, the overall transcriptional regulatory networks are highly divergent, implying that starch metabolism shifts from growth-oriented anabolism during development to stress-oriented homeostasis maintenance during storage.

Collectively, the stage-dependent transcriptional variation observed in this study illustrates a clear partition of physiological functions across tuber lifespan. Growth-stage metabolism focuses on efficient starch synthesis and accumulation with conserved basal transcriptional regulation, while storage-stage metabolism prioritizes stress resistance and starch stabilization with intensive genetic reprogramming. These temporal transcriptional characteristics provide a fundamental basis for dissecting stage-specific core genes and molecular pathways governing potato tuber starch homeostasis.

### 4.2 Correlation Analysis between GO/KEGG Enriched Pathways and Starch Metabolism

Integrated functional enrichment profiling indicated that cell wall remodeling modules, multi-stage metabolic pathways, and transcription factor regulatory networks act synergistically to mediate dynamic starch anabolism, catabolism, and homeostasis throughout potato tuber development and postharvest storage. Rather than functioning as independent regulatory units, these spatiotemporal modules form a coordinated regulatory system that adapts starch metabolic patterns to the physiological demands of tuber growth, maturation, and stress resistance.

#### 4.2.1 Functional implication of GO-based cell wall remodeling in starch metabolic regulation

XET-dominated cell wall remodeling represents a vital auxiliary regulatory mechanism underlying tuber starch metabolism, exerting stage-specific functional effects to support orderly starch turnover. Previous studies have confirmed that XET activity is essential for plant tissue morphogenesis and microenvironmental homeostasis, which indirectly shapes carbohydrate allocation and metabolic stability (Xiong *et al*., 2026; Yuan *et al*., 2024; Seale, 2020).

During tuber bulking, cell wall relaxation and structural reconstruction driven by XET activity optimizes source–sink transport efficiency. The modified cellular spatial structure facilitates the translocation of photosynthates to developing tubers, ensuring sufficient carbon skeleton supply for continuous starch synthesis and accumulation. This structural regulation mechanism complements the core starch anabolic pathway, revealing that cell wall plasticity is a prerequisite for efficient starch deposition during rapid tuber expansion (Hidvégi *et al*., 2024; Gong *et al*., 2025).

At the maturation stage, sustained cell wall remodeling primarily functions in metabolic stabilization rather than structural growth. Intact cell wall architecture shields intracellular starch metabolic enzymes from oxidative damage caused by senescence-related ROS accumulation, alleviating starch oxidative degradation. This protective mechanism sustains steady-state starch synthesis and locks peak starch reserves, which matches the stable storage physiological characteristics of mature tubers (Shirani-Bidabadi *et al*., 2024).

Under low-temperature storage conditions, cell wall polysaccharide remodeling serves as a core adaptive strategy linking stress response and starch homeostasis. Dynamic reconstruction of cell wall polysaccharide networks reshapes intracellular soluble carbon pools and coordinates with vitamin B6 metabolism and defense signaling. The enhanced cellular structural stability effectively resists cold-induced metabolic disorder, restricts excessive starch hydrolysis, and maintains long-term starch quality stability during storage (Zhang *et al*., 2017; Wang *et al*., 2026; Zhou *et al*., 2020).

#### 4.2.2 Mechanistic interpretation of KEGG metabolic pathways underlying stage-dependent starch homeostasis

Distinct KEGG-enriched pathways dominate starch metabolic regulation at different stages, reflecting the adaptive reprogramming of tuber carbon metabolism in response to developmental transitions and environmental changes.

In the bulking stage, photosynthetic carbon fixation, redox balance, and circadian rhythm signaling constitute the fundamental metabolic guarantee for rapid starch accumulation. Enhanced photosynthetic efficiency sustains continuous carbon assimilation and sucrose supply, providing material basis for starch biosynthesis (Geigenberger *et al*., 2004; Zierer *et al*., 2021).

Glutathione-mediated ROS scavenging stabilizes the activity of starch synthases, avoiding metabolic perturbation under vigorous cell proliferation (Shi *et al*., 2025). Meanwhile, circadian rhythm signaling temporally calibrates carbon allocation rhythms, optimizing the efficiency of starch anabolism and preventing invalid carbon consumption (Niu *et al*., 2022).

During tuber maturation, secondary metabolism and lipid metabolism drive the functional transition of tubers from rapid starch accumulation to stable storage status. Phenylpropanoid metabolism-derived lignin and phenolic compounds provide physical and antioxidant dual protection for starch granules, inhibiting premature starch degradation and improving storage stability (Navarre *et al*., 2013; Zhang *et al*., 2023). Continuous lipid synthesis maintains plastid membrane integrity, achieving spatial isolation between starch granules and hydrolytic enzymes. The physiological decline of photosynthetic carbon input actively terminates rapid starch accumulation, enabling tubers to enter a metabolically stable mature state (Geigenberger *et al*., 2004).

For postharvest cold storage, plant–pathogen interaction signaling and vitamin B6 metabolism are pivotal for maintaining starch homeostasis under abiotic stress. Cold-triggered calcium and ROS signaling cascades rewire carbohydrate metabolic flux, balancing starch synthesis and degradation to adapt to low-temperature stress (Fenyk *et al*., 2015; Wu *et al*., 2024). Vitamin B6 metabolism further consolidates starch stability by regulating starch hydrolase activity and eliminating oxidative damage to metabolic proteins, serving as a key protective pathway for starch quality maintenance during long-term storage (Mooney *et al*., 2013; Samsatly *et al*., 2020; Nasr *et al*., 2025).

#### 4.2.3 Transcriptional regulation of WRKY factors on starch homeostasis during storage

WRKY family transcription factors function as core transcriptional switches governing starch metabolic reprogramming during tuber cold storage, supplementing the structural and metabolic regulatory modules identified above. Multiple cross-species and omics studies have verified that WRKY proteins target conserved W-box motifs in starch metabolic gene promoters, serving as essential mediators linking stress signaling and carbon metabolism (Sun *et al*., 2003; Van Harsselaar *et al*., 2017).

Functional evidence has demonstrated that WRKY transcription factors exert dual regulatory effects on starch turnover, activating anabolic pathways while repressing catabolic processes to balance starch and soluble sugar ratios under stress conditions (Chiab, 2021). Combined with the reported universal regulatory role of potato WRKY members in developmental starch flux modulation, the identified StWRKY8, StWRKY9, and StWRKY22 are speculated to coordinate global starch transcriptional networks during storage. These TFs precisely fine-tune starch metabolic efficiency, offset cold stress-induced starch degradation, and sustain long-term starch homeostasis in postharvest tubers.

#### 4.2.4 Integrated regulatory network of tuber starch homeostasis

In conclusion, potato tuber starch homeostasis is governed by a multi-layered spatiotemporal regulatory network integrating structural modification, basal metabolism, and transcriptional regulation. During bulking, cell wall structural plasticity coupled with photosynthetic, redox and circadian signaling supports efficient starch accumulation. At maturation, secondary metabolism and lipid remodeling establish stable starch storage status via physical and antioxidant protection. During cold storage, cell wall carbon redistribution, stress-responsive metabolic pathways, and WRKY-mediated transcriptional modulation synergistically stabilize starch turnover. This stage-specific and multi-module coordinated mechanism elaborates the molecular basis of dynamic starch metabolism throughout tuber development and storage, providing new insights into the regulatory network of potato starch quality formation.

### 4.3 Functional Analysis of Key Stage-Persistent Genes in Starch and Sucrose Metabolism

Starch and sucrose metabolism represents the core functional pathway governing tuber starch anabolism and catabolism throughout development and storage. Combined with parental polymorphic SNP variation, population enzyme activity verification and qRT-PCR validation, two stage-persistent core genes (*Soltu.DM.02G019910* and *Soltu.DM.02G015310*) were identified as critical negative and positive regulators of tuber starch homeostasis, respectively. Their functional divergence and stage-specific regulatory effects largely account for the differential starch accumulation phenotypes between high– and low-starch potato materials.

#### 4.3.1 Negative regulatory effect of *Soltu.DM.02G019910* on starch accumulation

*Soltu.DM.02G019910* encodes β-glucosidase, a key enzyme responsible for cellulose and oligosaccharide hydrolysis to generate intracellular glucose. In general plant metabolic systems, glucose derived from cell wall polysaccharide hydrolysis acts as a direct carbon substrate to facilitate starch biosynthesis (Xiao *et al*., 2023; Zhang *et al*., 2026). However, the persistent high expression and elevated β-glucosidase activity in low-starch potato populations indicate a distinct negative regulatory mechanism in potato tubers. This inconsistent phenotype can be explained by threshold-dependent sugar feedback regulation and carbon resource redirection.

Excess glucose produced by continuously enhanced β-glucosidase activity fails to participate in starch synthesis. Instead, redundant carbon sources are preferentially consumed via cell wall reconstruction and respiratory metabolism, resulting in carbon diversion and insufficient carbon supply for starch deposition (Ferreira and Sonnewald, 2012). Furthermore, excessive intracellular glucose triggers conserved hexokinase-dependent sugar signaling feedback inhibition. High glucose levels suppress the activity of rate-limiting starch synthetic enzymes including AGPase, GBSS and SS, fundamentally blocking starch anabolic flux (Tiessen *et al*., 2002; McKibbin *et al*., 2006). Such feedback repression only occurs when glucose concentration exceeds the physiological threshold, which explains the discrepancy between the present results and previous reports that moderate glucose supply promotes starch synthesis. Combined with population enzyme activity verification, this evidence confirms that sustained high expression of *Soltu.DM.02G019910* restricts starch accumulation by disrupting carbon allocation balance and inhibiting core starch synthetic pathways, acting as a negative regulator of tuber starch accumulation.

#### 4.3.2 Positive regulatory effect of *Soltu.DM.02G015310* on starch homeostasis

*Soltu.DM.02G015310* participates in trehalose metabolism by regulating α,α-trehalose phosphatase and trehalose-6-phosphate phosphatase activity, which modulates the dynamic balance of intracellular trehalose-6-phosphate (T6P) and trehalose. Trehalose metabolism is a classic regulatory module balancing starch turnover and stress adaptation in sink organs (Li *et al*., 2022). T6P acts as a pivotal sugar signal that negatively regulates starch synthesis, whereas trehalose stabilizes starch structure and suppresses starch degradation, making the T6P/trehalose ratio a key determinant of starch homeostasis.

The stage-specific expression pattern of *Soltu.DM.02G015310* precisely matches the physiological rhythm of tuber starch metabolism. Its higher expression in high-starch tubers during bulking and storage stages accelerates the conversion of T6P to trehalose, reduces T6P accumulation, and relieves T6P-mediated inhibition of starch synthesis. Meanwhile, elevated trehalose content inhibits starch hydrolase activity and alleviates stress-induced starch degradation, thereby maintaining efficient starch accumulation and stable starch quality. During the maturation stage, relatively low gene expression enables moderate T6P accumulation, which stabilizes metabolic homeostasis and prevents excessive starch metabolic fluctuation, adapting to the physiological characteristics of peak starch storage (Debast *et al*., 2011). This stage-dependent regulatory mode enables *Soltu.DM.02G015310* to positively mediate starch synthesis and stabilization throughout tuber development and storage.

#### 4.3.3 Genetic basis and synergistic regulatory model of starch metabolic genes

Parental polymorphic SNP analysis further explained the intrinsic genetic variation underlying the differential expression of the two genes. The high polymorphism proportion of *Soltu.DM.02G019910* and *Soltu.DM.02G015310* indicates that their expression divergence between high– and low-starch populations is derived from parental genetic allelic variation. The high-starch parent contributes superior starch-favorable haplotypes, which repress *Soltu.DM.02G019910* expression and promote *Soltu.DM.02G015310* expression, ultimately forming a favorable genetic background for high starch accumulation. Collectively, the two stage-persistent genes form a dual antagonistic regulatory module governing tuber starch metabolism. *Soltu.DM.02G019910* mediates carbon diversion and glucose feedback inhibition to restrict starch accumulation, while *Soltu.DM.02G015310* modulates trehalose signaling to stabilize starch anabolism and inhibit starch degradation. The qRT-PCR and population enzyme activity validation further verified the reliability of transcriptome results and the biological function of key genes. This two-gene synergistic regulatory model complements the multi-layered starch metabolic network, elaborating the molecular and genetic basis of phenotypic differences in potato tuber starch content.

## 5. Conclusion

This study systematically explored the molecular and regulatory mechanisms of starch metabolism in potato tubers during bulking, maturation and storage stages via transcriptome profiling, functional enrichment and genetic variation analysis. It revealed prominent stage-specific transcriptional and metabolic regulatory features of tuber starch metabolism, with stable transcription sustaining starch synthesis in developmental stages and extensive transcriptomic reprogramming switching physiological functions for stress adaptation during storage. GO-based cell wall remodeling and temporally specific KEGG pathways synergistically control dynamic starch homeostasis across different stages, while 24 core starch and sucrose metabolism-related differentially expressed genes form stage-distinct metabolic patterns. Two persistently and antagonistically expressed core genes *Soltu.DM.02G019910* and *Soltu.DM.02G015310*, combined with differential enzyme activities and parental SNP genetic variations, underpin the starch phenotypic divergence among potato germplasms. Furthermore, the study constructed a multi-layered spatiotemporal regulatory network of potato starch metabolism, offering solid theoretical support and key genetic resources for high-quality, high-starch potato molecular breeding.

## Supplementary data

Table S1 Specific primers used for qRT-PCR amplification of key genes involved in potato starch and sucrose metabolism

Table S2 Summary of transcriptome sequencing quality statistics for potato tuber samp

Table S3 List of all expressed genes detected in the starch and sucrose metabolic pathways of potato tubers.

Table S4 Quality parameters of whole-genome DNA resequencing data for Huasheng No.7, Yunshu 306, and their hybrid progenies

Table S5 Starch content and β-glucosidase activity of potato breeding population grown under winter cultivation conditi

Fig. S1 Detailed information of advanced-generation progenies derived from the cross combination of Huasheng No.7.

Fig. S2 Schematic workflow of potato tuber sample collection.

Fig. S3 On-site photograph of harvested potato tuber samples.

Fig. S4 Correlation heatmaps of gene expression among all samples at three key developmental stages of potato tubers: tuber bulking stage (A), tuber maturation stage (B), and storage stage (C).

Fig. S5 Boxplots of gene expression (FPKM values) of individual samples at tuber bulking stage (A), tuber maturation stage (B), and storage stage (C).

Fig. S6 Enriched KEGG pathways of differentially expressed genes at the tuber bulking stage.

Fig. S7 Enriched KEGG pathways of differentially expressed genes at the tuber maturation stage.

Fig. S8 Enriched KEGG pathways of differentially expressed genes at the storage stage.

## Author contributions

YC: performed most of the experiments, data analysis, figure visualization and manuscript drafting; YX: provided experimental materials and research funding; ZL: conducted raw data analysis of DNA resequencing; BJ: performed enzyme activity determination; JB, LZ, LH, AX, WJ and QP: assisted with sample collection and partial experimental work; SY and LJ: conceptualized and designed the study, revised the manuscript, supervised the research and administrated the project. All authors have read and approved the final manuscript.

## Conflict of interest

The authors have no conflict of interest to declare.

## Funding

This work was supported by Science and Technology Plan of the Inner Mongolia Autonomous Region (2024KJHZ0010).

## Data availability

All data are available by contacting with the corresponding author.

## Notes

### Competing Interest Statement

The authors have declared no competing interest.

## References

1. Abeuova L, Kali B, Tussipkan D, Akhmetollayeva A, Ramankulov Y, Manabayeva S. 2023. CRISPR/Cas9-mediated multiple guide RNA-targeted mutagenesis in the potato. Transgenic Research, 32, 383–397.

2. Adegbaju MS, Gouws N, van der Vyver C, Claassens P, Kossmann J, Fischer Stettler M, Lloyd JR. 2025. Simultaneous Repression of GLUCAN WATER DIKINASE 1 and STARCH BRANCHING ENZYME 1 in Potato Tubers Leads to Starch With Increased Amylose and Novel Industrial Properties. Biotechnology Journal, 20, e70051.

3. Ahmad D, Zhang L, Gao Y, Deng B, Wu D, Bao J. 2026. Genotypic diversity and genome-wide association study of gelatinization and retrogradation properties of potato starch. Food Chemistry: Molecular Sciences, 100396.

4. Ai J, Yang M, Zou J, Wu Y, Li C, Wang Y, Gao D. 2025. The transcription factor StERF75 negatively regulates starch biosynthesis by targeting Isoamylase in potato. International Journal of Biological Macromolecules, 146118.

5. Ali NM, Rashed MAE, Atta AH, Abd-Elhalim HM, Ahmed NE, Metry EA. 2023. Increasing of amylopectin starch content in tetraploid potato (Solanum tuberosum L.) using CRISPR/Cas9 system.

6. Bai JM, Yang QF, Yang WL, Li XP, Li SF, Yang LY, Sui QJ. 2006. Relationships of potato tuber yield and starch yield with varieties and cultivation practices. Chinese Potato Journal, 20, 160–162.

7. Cairns JRK, Esen, A. (010. β-Glucosidases. Cellular and molecular life sciences: CMLS, 67, 3389.

8. Carpenter M A., Joyce NI., Genet RA., Cooper RD., Murray SR., Noble AD, Timmerman-Vaughan GM. 2015. Starch phosphorylation in potato tubers is influenced by allelic variation in the genes encoding glucan water dikinase, starch branching enzymes I and II, and starch synthase III. Frontiers in Plant Science, 6, 143.

9. Chiab N. 2021. The Overexpression of The VvWRKY2 Transcription Factor in Potato Improved the Agricultural Performance and Tubers’ Physio-Chemical and Industrial Properties Even Under Non Stress Conditions.

10. Cui G, Zhou T, Liu Z, Wang T, Wang Q, Liu T. 202). Deciphering the regulatory mechanisms of potato cold-induced sweetening via integrated time-course transcriptome and metabolome analysis. Frontiers in Plant Science, 16, 1551265.

11. Cui GC 2025. Molecular Mechanism of Heat Shock TranscriptionFactor StHSFA2 Regulating Cold-inducedSweetening in Potato [Master’s thesis, Shandong Agricultural University]. 10.27277/d.cnki.gsdnu.2025.000605

12. Debast S, Nunes-Nesi A, Hajirezaei MR, Hofmann J, Sonnewald U, Börnke F. 2011. Altering trehalose-6-phosphate content in transgenic potato tubers affects tuber growth and alters responsiveness to hormones during sprouting. Plant physiology, 156, 1754–1771.

13. Doyle JJ, Doyle JL. 1987. A rapid DNA isolation procedure for small quantities of fresh leaf tissue. Phytochemical bulletin.

14. Fan SH, Dong QS, Zhao TJ, Xie GQ, Wang Y, Zhao YT, Shi XR. 2019. Development and application of HRM molecular markers for high amylopectin potato mutant ‘P24023’. Chinese Potato Journal, 33(6), 338–343. Chinese Potato Journal, 33, 338–343.

15. Fenyk S, Townsend PD, Dixon CH, Spies GB, Campillo ADSE, Cann MJ. 2015. The potato nucleotide-binding leucine-rich repeat (NLR) immune receptor Rx1 is a pathogen-dependent DNA-deforming protein. Journal of Biological Chemistry, 290, 24945–24960.

16. Ferreira SJ, Sonnewald U. 2012. The mode of sucrose degradation in potato tubers determines the fate of assimilate utilization. Frontiers in plant science, 3, 23.

17. Geigenberger P, Stitt M, Fernie AR. 2004. Metabolic control analysis and regulation of the conversion of sucrose to starch in growing potato tubers. Plant, Cell & Environment, 27, 655–673.

18. Gong H, Li H, Wang C, Kui Q, Dusengemungu L, Cai X, Feng Z. 2025. StSUT2 Regulates Cell Wall Architecture and Biotic Stress Responses in Potatoes (Solanum tuberosum). Plants, 14, 2941.

19. Gong X, Zhou Y, Qin Q, Wang B, Wang L, Jin C, Fang W. 2024. Nitrate assimilation compensates for cell wall biosynthesis in the absence of Aspergillus fumigatus phosphoglucose isomerase. Applied and Environmental Microbiology, 90, e01138–24.

20. Haase NU. 2003. Estimation of dry matter and starch concentration in potatoes by determination of under-water weight and near infrared spectroscopy. Potato research, 46, 117–127.

21. Harris HC, Warren FJ. 2024. The impact of Cas9-mediated mutagenesis of genes encoding potato starch-branching enzymes on starch structural properties and in vitro digestibility. Carbohydrate Polymers, 345, 122561.

22. Hidvégi N, Dobránszki J, Tóth B, Gulyás A. 2024. Expression responses of XTH genes in tomato and potato to environmental mechanical forces: focus on behavior in response to rainfall, wind and touch. Plant Signaling & Behavior, 19, 2360296.

23. Hochmuth A, Carswell M, Rowland A, Scarbrough D, Esch L, Kamble NU, Seung D. 2025. Distinct effects of PTST2b and MRC on starch granule morphogenesis in potato tubers. Plant Biotechnology Journal, 23, 412–429.

24. Hu S, Song L, Yi X, Ni X, Cui X, Duan S, Liu X. 2025. Potato STARCH SYNTHEASE 5 is critical for simple starch granule initiation in amyloplasts and tuber development. The Plant Journal, 122, e70206.

25. Jayarathna S, Hofvander P, Péter-Szabó Z, Andersson M, Andersson R. 2024. GBSS mutations in an SBE mutated background restore the potato starch granule morphology and produce ordered granules despite differences to native molecular structure. Carbohydrate Polymers, 331, 121860.

26. Jiang B, Ao X, Yu XG, Liu ZR, Li H, Wang GP, Ren K, Tang CS. 2024. A New Starch Processing Potato Variety ‘Huasheng 7’. Chinese Potato Journal, 38, 206–208

27. Jiang B, Ren K, Wang GP. 2020. Multi-resistant Starch Processing Potato Variety ‘Villas’. Chinese Potato Journal 34, 191–192.

28. Jiang R, Jiang C, Ni X, Hu S, Cui X, Liu, X. 2025. Overexpression of the cold-repressed StWRKY70 enhances cold-induced sweetening resistance by coordinately attenuating starch-to-sugar conversion and increasing respiratory pathways. Postharvest Biology and Technology, 230, 113833.

29. Jin XH, Yu Z, Zhang X, Yu XX, Li JQ, Li JW. 2023. Development of SSR Molecular Markers and Preliminary Identification of Candidate Gene Associated with Potato Starch Content. Molecular Plant Breeding, 21, 2671–2676.

30. Khan S, Korai Z, Yang L, Korai SK, Li S, Wang X. 2026. Metabolic reprogramming in plant defense: linking signaling networks to metabolomics-driven insights. Plant Signaling & Behavior, 21, 2672221.

31. Khlestkin VK, Erst TV, Rozanova IV, Efimov VM, Khlestkina EK. 2020. Genetic loci determining potato starch yield and granule morphology revealed by genome-wide association study (GWAS). PeerJ, 8, e10286.

32. Khlestkin VK, Rozanova IV, Efimov VM, Khlestkina EK. 2019. Starch phosphorylation associated SNPs found by genome-wide association studies in the potato (Solanum tuberosum L.). BMC genetics, 20, 29.

33. Kolbe A, Tiessen A, Schluepmann H, Paul M, Ulrich S, Geigenberger P. 2005. Trehalose 6-phosphate regulates starch synthesis via posttranslational redox activation of ADP-glucose pyrophosphorylase. Proceedings of the National Academy of Sciences, 102, 11118–11123.

34. Kumar S, Bandyopadhyay N, Saxena S, Hajare SN, More V, Gautam S. 2024. Differential gene expression in irradiated potato tubers contributed to sprout inhibition and quality retention during a commercial scale storage. Scientific Reports, 14, 13484.

35. Larichev K, Sergeeva E, Karetnikov D, Salina E, Kochetov A. 2022. Development of CRISPR/Cas9-based genome editing constructs for Nevsky and Udacha potato cultivars. In Bioinformatics of Genome Regulation and Structure/Systems Biology (BGRS/SB-2022) (pp. 621–621).

36. Li C, Sun L, Zhu J, Ji X, Huang R, Ge Y. 2022. Trehalose regulates starch, sorbitol, and energy metabolism to enhance tolerance to blue mold of “Golden Delicious” apple fruit. Journal of agricultural and food chemistry, 70, 5658–5667.

37. Li HP, Liang X, Shen XS, Wang P. 2017. Breeding and Main Points of Cultivation Technique of the New High Starch and Early Maturing Potato Variety Chuanyu 16. Tillage and Cultivation, 37, 81–82.

38. Li J, Yu X, Zhang S, Yu Z, Li J, Jin X, Yang D. 2021. Identification of starch candidate genes using SLAF-seq and BSA strategies and development of related SNP-CAPS markers in tetraploid potato. PLoS One, 16, e0261403.

39. Li JW, Wen GH, Li GF, Wang YH, Jia XX, Zhang R, Ma S. 2020. Evaluation of Starch Yield and Nutrition Quality Traits in Longshu Series Potato Varieties With High Starch Content. Journal of Nuclear Agricultural Sciences, 34, 329–338.

40. Li L, Tacke E, Hofferbert HR, Lübeck J, Strahwald J, Draffehn AM, Gebhardt C. 2013. Validation of candidate gene markers for marker-assisted selection of potato cultivars with improved tuber quality. Theoretical and Applied Genetics, 126, 1039–1052.

41. Li YS, Xu NS, Sui QJ. 2018. A New High Starch Potato Variety-’Yunshu 202’. Chinese Potato Journal, 32, 124–126.

42. Li ZY. 2024. Function characterization of StNAC033 regulatingcarbon source allocation in potato [Master’s thesis, Southwest University]. 10.27684/d.cnki.gxndx.2024.004931

43. Liu SH. 1999. Breeding and use of potato with high-starch content — Neishu 7. Proceedings of the Annual Conference of Potato Professional Committee, Chinese Society of Crop Sciences, 54–56.

44. Liu T, Kawochar MA, Begum S, Wang E, Zhou T, Jing S, Song B. 2023. Potato tonoplast sugar transporter 1 controls tuber sugar accumulation during postharvest cold storage. Horticulture Research, 10, uhad035.

45. Liu T, Kawochar MA, Liu S, Cheng Y, Begum S, Wang E, Song B. 2023. Suppression of the tonoplast sugar transporter, StTST3. 1, affects transitory starch turnover and plant growth in potato. The Plant Journal, 113, 342–356.

46. Liu TT. 2023. The Function and Molecular Mechanism of ABA-MediatedStarch and Sugar Metabolism in Potato Freezing Tolerance [Master’s thesis, Huazhong Agricultural University]. 10.27158/d.cnki.ghznu.2023.002005

47. Locquet C, Berti M, Bray F, Hulin H, Stoclet G, Seung D, Szydlowski N. 2026. LIKE EARLY STARVATION is involved in the regulation of starch initiation in potato tubers (Solanum tuberosum cv. Désirée). Journal of Experimental Botany, 77, 1789–1806.

48. Ma L, Liu Y, Han Y, Deng H, Jiang H, Prusky, D. 2023. Mechanical wounds expedited starch degradation in the wound tissues of potato tubers. International Journal of Biological Macromolecules, 236, 124036.

49. Ma YY, Guo ZM, Zhang X, Li ZB, Li K, Guo HC. 2025. Starch accumulation characteristics and expression of related synthase genes in tubers of different potato varieties. Journal of Northeast Agricultural University, 56, 12–22.

50. Manna B, Ghosh, A. 2021. Molecular insight into glucose-induced conformational change to investigate uncompetitive inhibition of GH1 β-glucosidase. ACS Sustainable Chemistry & Engineering, 9, 1613–1624.

51. McKibbin R S, Muttucumaru N, Paul MJ, Powers SJ, Burrell MM, Halford NG. 2006. Production of high starch, low glucose potatoes through over expression of the metabolic regulator SnRK1. Plant biotechnology journal, 4, 409–418.

52. Mooney S, Chen L, Kühn C, Navarre R, Knowles N R, Hellmann H. 2013. Genotype specific changes in vitamin B6 content and the PDX family in potato. BioMed research international, 2013, 389723.

53. Naher J, Sabuj ZH, Sumona SI, Chakraborty SP, Hossain MR, Nath UK. 2024. Heat stress modulates superoxide and hydrogen peroxide dismutation and starch synthesis during tuber development in potato. American Journal of Potato Research, 101, 275–289.

54. Nasr Esfahani M, Koch L, Hofmann J, Sonnewald S, Sonnewald U. 2025. Organ specific transcriptional and metabolic adaptations of potato plants to limited phosphate availability prior and after tuberization. The Plant Journal, 123, e70445.

55. Navarre DA, Payyavula RS, Shakya R, Knowles NR, Pillai SS. 2013. Changes in potato phenylpropanoid metabolism during tuber development. Plant Physiology and Biochemistry, 65, 89–101.

56. Nicol F, His I, Jauneau A, Vernhettes S, Canut H, Höfte H. 1998. A plasma membrane bound putative endo 1, 4 β d glucanase is required for normal wall assembly and cell elongation in Arabidopsis. The EMBO journal, 17, 5563–5576

57. Niu Y, Li G, Jian Y, Duan S, Liu J, Jin L. 2022. Genes related to circadian rhythm are involved in regulating tuberization time in potato. Horticultural Plant Journal, 8, 369–380.

58. Richter C, Fulgoni K, Fulgoni III VL, Johnson B, Kijek M, Maniscalco S, Psota T. 2025. Assessment of the unique nutrient contribution of white potatoes in the diet and the nutrient implications of replacing Refined and Whole Grains with Starchy Vegetables. Frontiers in Nutrition, 12, 1692564.

59. Rodríguez-Saavedra C, Morgado-Martínez LE, Burgos-Palacios A, King-Díaz B, López-Coria M, Sánchez-Nieto S. 2021. Moonlighting proteins: the case of the hexokinases. Frontiers in molecular biosciences, 8, 701975.

60. Samsatly J, Bayen S, Jabaji SH. 2020. Vitamin B6 is under a tight balance during disease development by Rhizoctonia solani on different cultivars of potato and on Arabidopsis thaliana mutants. Frontiers in Plant Science, 11, 875.

61. Schönhals EM, Ortega F, Barandalla L, Aragones A, Ruiz de Galarreta JI, Liao JC, Gebhardt C. 2016. Identification and reproducibility of diagnostic DNA markers for tuber starch and yield optimization in a novel association mapping population of potato (Solanum tuberosum L.). Theoretical and Applied Genetics, 129, 767–785.

62. Schreiber L, Nader-Nieto AC, Schönhals EM, Walkemeier B, Gebhardt C. 2014. SNPs in genes functional in starch-sugar interconversion associate with natural variation of tuber starch and sugar content of potato (Solanum tuberosum L.). G3: Genes, Genomes, Genetics, 4, 1797–1811.

63. Seale M. 2020. Cell wall remodeling during wood development. Plant Physiology, 182, 1800–1801

64. Sergeeva EM, Larichev KT, Salina EA, Kochetov AV. 2022. 32Starch metabolism in potato Solanum tuberosum L. Vavilov Journal of Genetics and Breeding, 26, 250.

65. Seung D, Schreier TB, Bürgy L, Eicke S, Zeeman SC. 2018. Two plastidial coiled-coil proteins are essential for normal starch granule initiation in Arabidopsis. The Plant Cell, 30, 1523–1542.

66. Seung D, Soyk S, Coiro M, Maier BA, Eicke S, Zeeman SC. 2015. PROTEIN TARGETING TO STARCH is required for localising GRANULE-BOUND STARCH SYNTHASE to starch granules and for normal amylose synthesis in Arabidopsis. PLoS biology, 13, e1002080.

67. Sharma S, Friberg M, Vogel P, Turesson H, Olsson N, Andersson M, Hofvander P. 2023. Pho1a (plastid starch phosphorylase) is duplicated and essential for normal starch granule phenotype in tubers of Solanum tuberosum L. Frontiers in Plant Science, 14, 1220973.

68. Shi N, Fan Y, Zhang W, Zhang Z, Pu Z, Sun C. 2025. Genome-wide identification and drought-responsive functional analysis of the GST gene family in potato (Solanum tuberosum L.). Antioxidants, 14, 239.

69. Shi Q, Xia Y, Xue N, Wang Q, Tao Q, Li G. 2024. Modulation of starch synthesis in Arabidopsis via phytochrome B mediated light signal transduction. Journal of Integrative Plant Biology, 66, 973–985.

70. Shirani-Bidabadi M, Nazarian-Firouzabadi F, Sorkheh K, Ismaili A. 2024. Transcriptomic analysis of potato (Solanum tuberosum L.) tuber development reveals new insights into starch biosynthesis. PLoS One, 19, e0297334.

71. Singh A, Compart J, AL Rawi SA, Mahto H, Ahmad AM, Fettke J. 2022. LIKE EARLY STARVATION 1 alters the glucan structures at the starch granule surface and thereby influences the action of both starch synthesizing and starch degrading enzymes. The Plant Journal, 111, 819–835.

72. Sowokinos JR, Preiss J. 1982. Pyrophosphorylases in Solanum tuberosum: III. purification, physical, and catalytic properties of ADPglucose pyrophosphorylase in potatoes. Plant Physiology, 69, 1459–1466.

73. Stein O, Granot D. 2019. An overview of sucrose synthases in plants. Frontiers in plant science, 10, 95.

74. Sun C, Palmqvist S, Olsson H, Borén M, Ahlandsberg S, Jansson C. 2003. A novel WRKY transcription factor, SUSIBA2, participates in sugar signaling in barley by binding to the sugar-responsive elements of the iso1 promoter. The Plant Cell, 15, 2076–2092.

75. Sun H. 2023. Function of StWRKY23 transcription factor inpotato tuber development and starchmetabolism of potato. (Master’s thesis, Southwest University). 10.27684/d.cnki.gxndx.2023.004692

76. Tiessen A, Hendriks JH, Stitt M, Branscheid A, Gibon Y, Geigenberger P. 2002. Starch synthesis in potato tubers is regulated by post-translational redox modification of ADP-glucose pyrophosphorylase: a novel regulatory mechanism linking starch synthesis to the sucrose supply. The Plant Cell, 14, 2191–2213.

77. Van Harsselaar JK, Lorenz J, Senning M, Sonnewald U, Sonnewald S. 2017. Genome-wide analysis of starch metabolism genes in potato (Solanum tuberosum L.). BMC genomics, 18, 37.

78. Wang W, Wang Z, Wei Y, Shang M, Ma R, He S. 2026. Natural allelic variations in IbJAZ10 IbNF YA3 complex regulate Rhizopus soft rot resistance in sweet potato. New Phytologist.

79. Waterschoot, J., Gomand, S. V., Fierens, E., & Delcour, J. A. (2015). Production, structure, physicochemical and functional properties of maize, cassava, wheat, potato and rice starches. Starch Stärke, 67(1-2), 14–29.

80. Wiberley-Bradford AE, Busse JS, Jiang J, Bethke PC. (014. Sugar metabolism, chip color, invertase activity, and gene expression during long-term cold storage of potato (Solanum tuberosum) tubers from wild-type and vacuolar invertase silencing lines of Katahdin. BMC Research Notes, 7, 801.

81. Wu C, Fang XT, Shi XX, Zhou X, Luo C, Zheng SL. 2024. Research Progress and Prospects on Cold-induced Sweetening in Potato Tubers. Chinese Potato Journal, 38, 168.

82. Wu TM, Lin WR, Kao CH, Hong CY. 2015. Gene knockout of glutathione reductase 3 results in increased sensitivity to salt stress in rice. Plant Molecular Biology, 87, 555–564.

83. Xiao Y, Dong S, Liu YJ, You C, Feng Y, Cui Q. 2023. Key roles of β-glucosidase BglA for the catabolism of both laminaribiose and cellobiose in the lignocellulolytic bacterium Clostridium thermocellum. International Journal of Biological Macromolecules, 250, 126226.

84. Xiong M, Zhang T, Qian X, Kadeer A, Kurotani K I, Huang Y. 2026. Xyloglucan endotransglucosylase/hydrolase family genes are required for the plant graft union formation through callus proliferation. Plant Physiology, 200, kiag030.

85. Xu X, Lian Y. 1997. Study on the development mechanism of potato tubers. Chinese Potato Journal, (2), 52–56.

86. Yang T, Li H, Li L, Wei W, Huang Y, Xiong F, Wei M. 2023. Genome-wide characterization and expression analysis of α-amylase and β-amylase genes underlying drought tolerance in cassava. BMC genomics, 24, 190.

87. Yuan W, Yao F, Liu Y, Xiao H, Sun S, Zhang J. 2024. Identification of the xyloglucan endotransglycosylase/hydrolase genes and the role of PagXTH12 in drought resistance in poplar. Forestry Research, 4, e039.

88. Zeeman SC, Kossmann J, Smith AM. 2010. Starch: its metabolism, evolution, and biotechnological modification in plants. Annual review of plant biology, 61, 209–234.

89. Zhang H, Hou J, Liu J, Zhang J, Song B, Xie C. 2017. The roles of starch metabolic pathways in the cold induced sweetening process in potatoes. Starch Stärke, 69, 1600194.

90. Zhang H, Zheng SY, Liang SX, Chen GF, Wang MY. 2019. Research Progress in Breeding Special Potatoes with High Starch Content. Crops, 35, 9–14.

91. Zhang J, Yao J, Mao L, Li Q, Wang L, Lin Q. 2023. Low temperature reduces potato wound formation by inhibiting phenylpropanoid metabolism and fatty acid biosynthesis. Frontiers in Plant Science, 13, 1109953.

92. Zhang Y, Chen M, Wang Y, Xiong Z, Xu K, Chen, H. 2026. High-efficiency enzymatic conversion of cellulose to starch by strengthening glucose-utilization bypass. Bioresource Technology, 134274.

93. Zhao CB, Song SY, Zhang CW, Wen T, Sun K. 2011. Analysis and Assessment on Nutritional Quality of Potato Varieties. Jilin Agricultural Sciences, 36, 58–60.

94. Zhao X, Jayarathna S, Turesson H, Fält AS, Nestor G, Gonzalez MN, Andersson M. 2021. Amylose starch with no detectable branching developed through DNA-free CRISPR-Cas9 mediated mutagenesis of two starch branching enzymes in potato. Scientific reports, 11, 4311.

95. Zhou W, Zhao S, He S, Ma Q, Lu X, Zhang P. 2020. Production of very high amylose cassava by post transcriptional silencing of branching enzyme genes. Journal of integrative plant biology, 62, 832–846.

96. Zhu X, Richael C, Chamberlain P, Busse JS, Bussan AJ, Jiang J, Bethke PC. 2014. Vacuolar invertase gene silencing in potato (Solanum tuberosum L.) improves processing quality by decreasing the frequency of sugar-end defects. PloS one, 9, e93381.

97. Zierer W, Rüscher D, Sonnewald U, Sonnewald S. 2021. Tuber and tuberous root development. Annual review of plant biology, 72, 551–580.

